# Cooperative mixing through hydrodynamic interactions in *Stylonychia lemnae*

**DOI:** 10.1101/2025.02.14.638298

**Authors:** Régis Turuban, Giovanni Noselli, Alfred Beran, Antonio DeSimone

## Abstract

Aquatic microorganisms typically inhabit a heterogeneous resource landscape, composed of localized and transient patches. To effectively exploit these resources, they have evolved a wide range of feeding strategies that combine chemotactic motility with active feeding flows. However, there is a notable lack of experimental studies that examine how these active flows shape resource fields to optimize feeding. In particular, the suspected cooperative hydrodynamics provided by groups of cells remains largely unexplored due to the difficulties in visualizing these dynamic three-dimensional flows. Here, we experimentally investigate how *Stylonychia lemnae* ciliates form feeding clusters of independent cells around food patches. Individual feeding flows interact hydrodynamically to create a chaotic collective flow at the population scale. Using a combination of experimental and numerical techniques, we measure and predict the entire collective flow, enabling us to assess its outstanding mixing and dispersion properties. We show that the active spreading of the food patch accelerates its detection by starving cells. As many additional fitness advantages provided by collective flows can be envisioned, we propose that this feeding cluster represents a form of intra-specific by-product cooperative behavior.

---

Survival for aquatic microorganisms is about encountering: maximizing interactions with prey or nutrients for feeding and minimizing encounters with predators. Often, interactions are mediated by the chemical signals constantly released by microorganisms, whether they wish it or not. To best exploit patchy resource fields (1–3) in a viscous world, microorganisms have evolved the ability to navigate concentration gradients toward the source (chemotaxis) and/or actively generate flows powerful enough to shape concentration gradients directly. Examples of such actively generated flows include the currents around the organism’s body as it swims (4–6), feeding flows produced by microorganisms either anchored to a solid substrate (7–11), tethered by gravity (12), or bonded together within a colony (13–16), as well as hydrodynamics waves triggered by rapid body contractions (17) or the long-range coherent flows produced by active suspension of pushers (18–20). Faced with this diversity of cell organizations and their associated active flows, scientists have long wondered which fluid kinematics would lead to optimal feeding rates/nutrient uptake, and under which specific conditions.

Particularly well-suited to tackle this problem are the organisms exhibiting different morphotypes, ranging from unicellular to undifferentiated multicellular colonies. For example, (13, 21) have shown that *volvox* colonies - spherical colonies made up of thousands of Chlamydomonas cells bonded to- gether - balance their expanding metabolic needs as their radius *R* grows by generating flows of increasing magnitude that steepen concentration gradients and thus ultimately enhance the diffusion flux, scaling just as the expanding surface *Q* ∼ *R*^2^. Other research by (14) estimates that cooperative hydrodynamics within choanoflagellate colonies (which are attached together in the fluid bulk) increase the fresh fluid supply per cell compared to freely-swimming single cells. However, another study by (15) presents a contrasting view, predicting that choanoflagellate colonies are less advantageous in terms of feeding flux compared to single anchored cells, which in turn are less advantageous than freely swimming cells. Although the prediction that freely swimming provides the highest feeding rate is consistent with the results of (22), it contradicts experimental findings indicating that cells tend to promote swimming over anchoring and feeding when deprived of food (23). Moreover, the claim that colonies are the least advantageous organization is challenged by the findings of (24), which showed that in some choanoflagellate species, colony formation is triggered by sensing chemicals produced by their prey bacteria, even at very low concentrations. Clearly, the systematic feeding advantages conferred by diverse fluid kinematics - especially in the case of colonies - are still poorly understood.

To reconcile predictions with observations, a better understanding of the dynamic interaction between organisms and the nutrient/prey field is required. So far, the vast majority of studies have based their numerical estimations of feeding rates on strong simplifications: a cell generating a steady active flow, in a still ambient fluid, and immersed in a uniform resource field. The latter two conditions hold true only for Pe_ambient_ *<* 1, yet many organisms operate in the Pe_ambient_ *>* 1 regime (25) (where Pe_ambient_ = *v*_ambient_*L*/*D* is the Péclet number relative to the ambient velocity field *v*_ambient_, *L* the characteristic flow size, and *D* the diffusion coefficient). Regarding the first condition, unsteadiness is rarely discussed, although (26, 27) have demonstrated that the simple case of two nearby sessile cells alternating feeding currents can drastically enhance feeding by introducing Lagrangian chaos. In fact, experimental observations of microorganisms interacting with a resource field in the large Pe regime highlight much more dynamic behavior, as illustrated by the striking collective behavior exhibited by two bacteria species, *Thiovulum majus* and *Uronemella* (28–30). In this case, the authors suggest that the cell’s oxygen consumption results from the highly dynamic interplay between three components: (i) their locomotion behavior, which depends on gradients of both *O*_2_ and population density; (ii) the macroscopic active flow produced by the community, controlled by local population density; and (iii) the *O*_2_ concentration field advected by the macroscopic flow, and consumed by the cells. Thus, these studies underscore the need to revisit the feeding problem in a patchy food environment by taking into account this complex coupling. This requires experimental measurements of the evolving 3D active flow, which, to our knowledge, have not been performed yet.

Here we propose to experimentally investigate the feeding behavior of *Stylonychia lemnae* (31) immersed in a patchy food environment. *S. lemnae* is a 200 *µ*m long freshwater ciliate belonging to Spirotrichea, a class of ciliates widely distributed across diverse ecosystems, including freshwater, saltwater, soils, and activated sludge (32). *S. lemnae* can either swim or attach itself to interfaces and crawl. Being anchored to solid boundary allows the cell to exert a net force on the fluid using its adoral zone of membranelles (AZM, see Fig. 1A-B and movie 1-2), directing the fluid toward its buccal apparatus. Our research reveals that *S. lemnae* exhibits a collective feeding behavior that closely resembles that of *T. majus* and *Uronemella*, characterized by individual chemosensory locomotion to locate and remain near food patches, resulting in cell clustering around these patches. Next, multiple hydrodynamic interactions lead to flow at the population scale. However, unlike *volvox*, choanoflagellates colonies or the veils of *Uronemella*, where cells are bound together, the cells in *S. lemnae* feeding clusters remain strictly independent. This leads to an important question: Can such a loosely organized group of cells, lacking physical connections, exhibit cooperative behavior through hydrodynamic interactions? Addressing this question could yield key insights into the selective forces that promoted the early evolution of multicellular life. To this end, we experimentally characterize both locomotion behavior and the full 3D unsteady flow generated by a feeding cluster of *S. lemnae*. We numerically quantify the dispersion and stretching kinematics of the collective flow. By combining these two elements within the lamellar approach (33, 34), we are able to predict the rate at which any patchy resource field is homogenized within the feeding cluster.

**Fig. 1.**
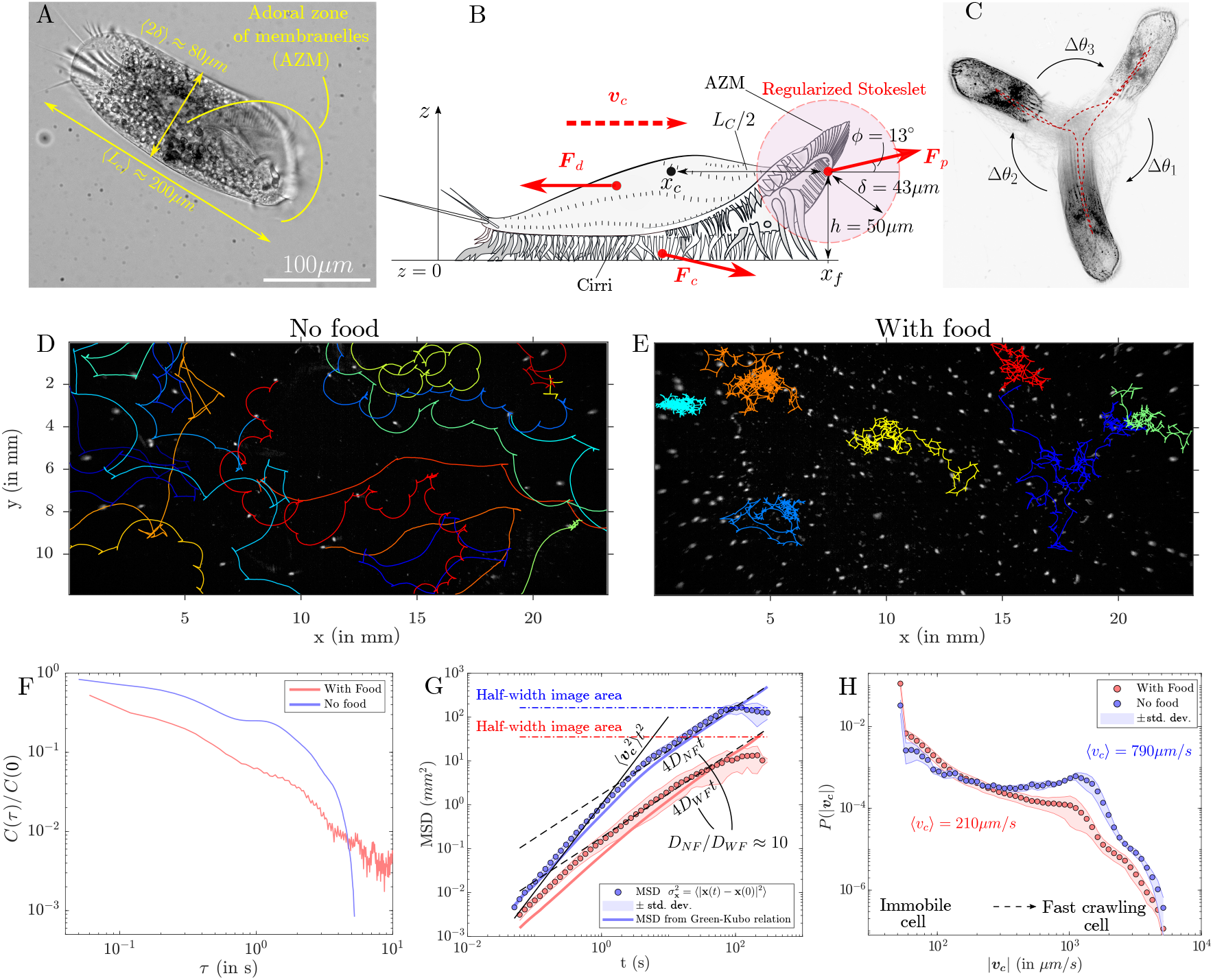
*S. lemnae*’s locomotion behavior. (A) Microscopy image of a *S. lemnae* cell viewed from below. (B) Schematic lateral view (adapted from the *Stylonychia* Wikipedia webpage), and the balance of forces acting on the cell body during crawling locomotion at speed *v*_*c*_. (C) A *S. lemnae* cell performing a series of three Side-Stepping Reactions (SSRs), a characteristic motion of the Spirotrich group. The cell’s trajectory is displayed as a red dashed line. (D) Trajectories ***x***_*i*_(*t*) followed by starving *S. lemnae* as they crawl on the bottom of their Petri dish culture. (E) Trajectories followed by the same cells ∼1-2 min after the injection and manual mixing of food. These trajectories are selected so that they do not overlap. (F) Normalized autocorrelation function of the crawling velocities ***v***_***c***_ (VAFs) for the two distinct ambient conditions: no food (in blue) and with food (in red). (G) Mean square displacement (MSD) calculated either from the variance of cell displacements (dots), or from the Green-Kubo relation (continuous lines), computed from the VAFs in panel (F). Same color code as before. Error bars (shaded area) indicate the standard deviation across seven different datasets. The continuous black line represents ballistic transport at early times, scaling as 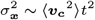, for the no-food case. The black dashed lines represent the asymptotic Fickian regime 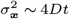 reached in both cases, with different dispersion coefficient *D*_NF_ (no food) and *D*_WF_ (with food). The colored dashed lines represent the characteristic area limit of the image beyond which cells are no longer visible. (H) Probability density function of the Lagrangian velocities magnitude *P* (|***v***_***c***_|) for the two ambient conditions. The minimum measurable velocity with our setup is 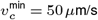, below which the cell is considered immobile.

## 1. Sensing food triggers a change of locomotion behavior in *S. lemnae*

In laboratory conditions, when starving, *S. lemnae* typically wanders on the floor of a Petri dish looking for food, as illustrated in Fig. 1D. Thanks to its cirri, it crawls in wide leftward arcs over distances up to several millimeters, with an average speed of about 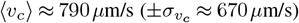, peaking at *v*_*c*_ ∼ 7000 *µ*m/s (i.e. 35*L*_*c*_/s, with *L*_*c*_ ≈ 200 *µ*m the average cell length). The leftward arcs are interspersed with sudden stops (*v*_*c*_ = 0), which can last from a few tens of milliseconds to a few tens of seconds. Then, either the cell resumes its movement forward, or it performs abrupt moves called Side-Stepping Reaction (SSR), typically observed in Spirotrich species, such as *S. mytilus* or *Euplotes* (35–37). These movements can be decomposed into two steps: first bouncing backward over a short distance, then rotating clockwise by a varying angle Δ*θ* (with a maximal angular velocity of about 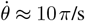). An example of a cell performing three successive SSR moves can be seen in Fig. 1C, resulting in that particular case in an almost complete revolution (movie 3).

We noticed a drastic change in locomotion behavior whenever *S. lemnae* senses food. To quantify it, we conducted the following experiment. In the first step, the positions of starving cells are tracked for 5 minutes. In the second step, we inject food (either yeast or *Chlorogonium* algae) and gently stir it by hand until the food concentration is homogeneous inside the whole Petri dish. Another sequence of images is then taken for 5 minutes. The cell trajectories are reconstructed using particle tracking velocimetry (PTV) and the Lagrangian velocities *v*_*c*_ extracted from the displacement between two successive images (*SI Appendix*, section 2A for the whole procedure). A first qualitative difference of locomotion patterns can be inferred from the comparison between Fig. 1D and Fig. 1E (movie 4). Clearly, cells deprived of food travel much farther whereas cells in the presence of food significantly reduce the length of their crawling segments in favor of prolonged stops. The cells still exhibit SSR movements, but because they travel less, the successive SSRs results in a rotation around a point, plus a small drift. At the macroscopic scale, the impact of this behavioral change is quantified through the mean squared displacement (MSD), defined as the variance of cells displacements 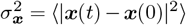, with ⟨…⟩ the ensemble average over all trajectories *i*, after checking that the cells spreading is isotropic, i.e. ⟨***v***_***c***_⟩ = 0. The MSDs in both cases are compared in Fig. 1H. The MSD for the no-food case well captures the early-time ballistic regime 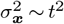. After a certain coherence time, the MSDs tends toward a diffusive regime 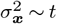, but this regime is soon constrained by the finite size limitation of our imaging setup (cells disappearing beyond the imaging area). To overcome this limitation and get the asymptotic scaling, we compute the MSD from the velocity autocorrelation function (VAF) (*SI Appendix*, section 2B), which are related through the expression 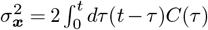,with *C*(*τ*) = ⟨***v***_***c***_(*τ*) · ***v***_***c***_(0)⟩, whose graphs are displayed on Fig. 1G, as a function of the time lag *τ* = *t*^′^ − *t*^′′^. The new MSD graphs are superimposed to the previous MSDs in the same Fig. 1H, and reveal that both MSDs reach an asymptotically Fickian regime 4*Dt* synonymous with Brownian-like motion, though with a dispersion coefficient *D* that varies with ambient conditions. These dispersion coefficients can be obtained through the Green-Kubo relation 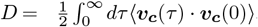. We get *D*_NF_ = 4.3 × 10^−7^ m^2^/s (no food) and *D*_WF_ = 4.5 × 10^−8^ m^2^/s (with food), indicating that cells searching for food span space *D*_NF_*/D*_WF_ ≈ 10 faster that cells cohabiting with food. The origin of the 10fold decrease of the dispersal rate is partly due to the slightly faster decay of *C*(*τ*) / *C*(0) as shown in Fig. 1F, but mainly due to the overall reduction in locomotion speed *v*_*c*_ through the ratio *C*(0)_NF_*/C*(0)_WF_ = ⟨***v***_***c***_^2^⟩ _NF_*/* ⟨***v***_***c***_^2^⟩ _WF_ ≈ 4.7. Indeed Fig. 1F displaying the distribution of Lagrangian velocities *P* (*v*_*c*_) shows how the food injection triggers a massive drop in the probability of high velocities, with the average velocity decreasing from ⟨*v*_*c*_⟩ ≈ 790 *µ*m/s down to ⟨*v*_*c*_⟩ ≈ 210 *µ*m/s. We conclude that the cells exhibit two distinct locomotion modes, switching from one to the other depending on environmental conditions, by relying significantly on the reduction of *v*_*c*_. As a matter of fact, modulating the locomotion speed to remain in a favorable area is one well-known chemotactic strategy (38).

## 2. Formation of a cluster of feeding cells around a food injection site and collective fluid mixing

To reproduce the patchy food environment in which many organisms live (1, 2), we conduct a similar experiment, with the difference that the food is now injected within a confined space (*SI Appendix*, section 3). A tiny amount of yeast (abundant on the cell scale) is injected onto the bottom plate among starving crawling cells, see Fig. 2A(i). The blob of food is originally *s*_0_ ≈ 0.5 mm = 2.5*L*_*c*_ in diameter. Before injection, cells exhibit persistent long traveling paths that, when integrated over a period of 5 s, frequently reach values up to 20*L*_*c*_ (with a maximum of 50*L*_*c*_). Immediately after injection, only the cells in close contact with the food injection point drastically reduce their traveled path. However, as time passes, the cells farther away also restrict their amplitude of motion, as showed by Fig. 2A(ii), *t* = 280 s after the food injection (movie 5). The transition towards shortened movement over time is indicated by the orange dashed line in Fig. 2B. After *t* = 400 s, the great majority of the cells visible in the image, even those far (up to 1 cm, i.e. 20*s*_0_) from the injection point, don’t travel: a feeding cell cluster has formed. The phenomenon is all the more intriguing given that yeast, a few micrometers wide, barely diffuses, with *D*_yeast_ ≈ 10^−11^ m^2^/s as given by the Stokes-Einstein equation. In addition, the formation of the cluster is reproducible systematically at each injection of yeast, regardless of the level of cell satiety, provided the environment is initially reasonably clear of food.

**Fig. 2.**
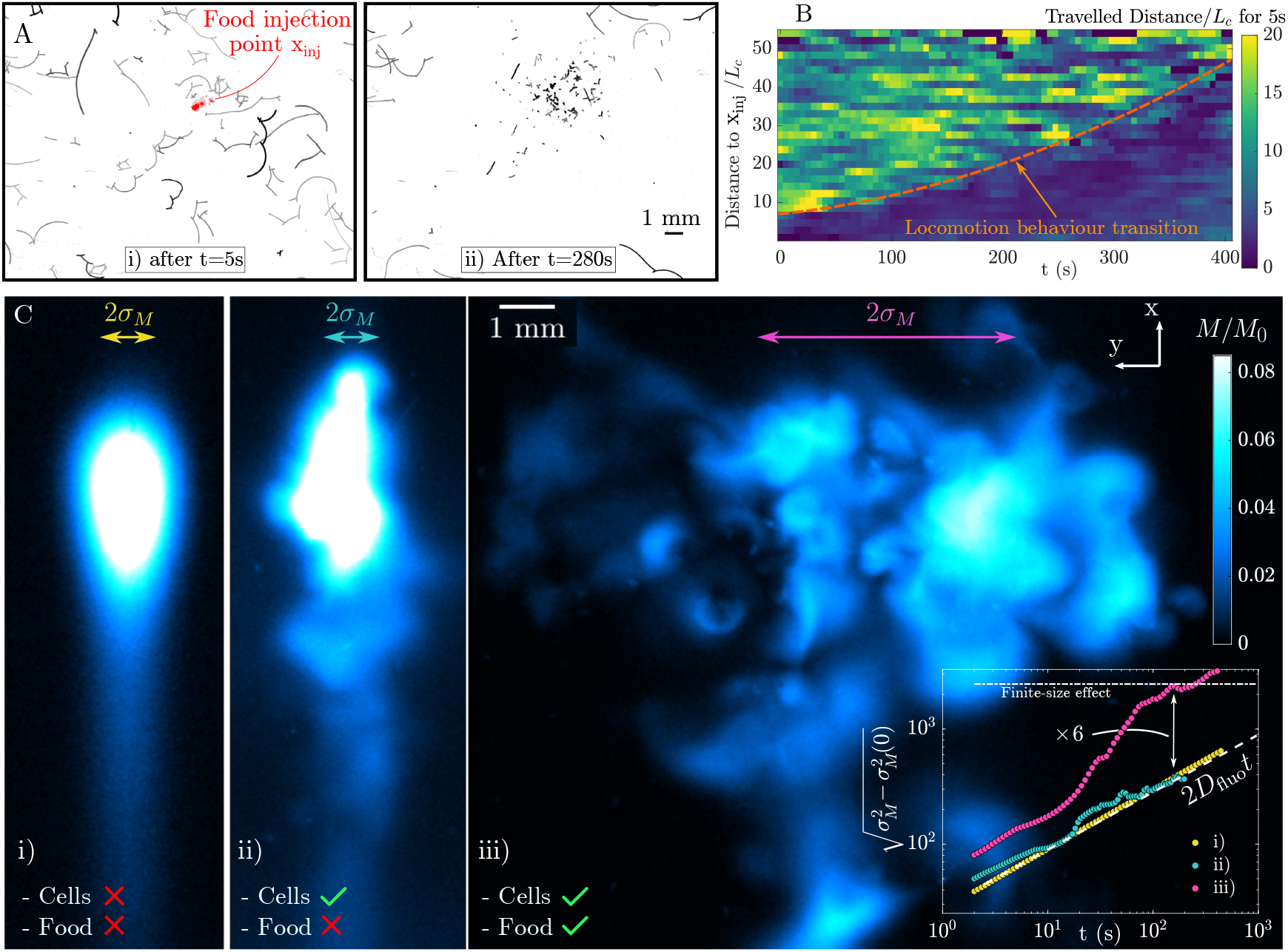
Collective flow by a feeding cell cluster. (A) Trajectories of a few *S. lemnae* integrated over 5 s, from i) *t* = 5 s and ii) *t* = 280 s after the local injection of yeast, at position x_inj_. (B) Temporal evolution of the distance traveled by different *S. lemnae* over 5 s, depending on their initial distance from the food injection point x_inj_. The statistical analysis is based on ≈ 640 trajectories from one experiment. The orange continuous line shows the transition between the two locomotion behavior, triggered further and further away from the food injection point as time progresses. Initially, about 50 cells are visible on the images, progressively increasing to reach up to 100 cells, 400 s after the injection. (C) Imaging of a fluorescein blob injected in the observation chamber either i) in the absence of *S. lemnae* cells, or ii) injected among cells, or iii) mixed with yeast prior to injection among cells. These snapshots were taken at *t* = 174 s after the injection of the dye blob. Fluorescence intensity levels *I*(*x, y*) are converted to mass levels *M* (*x, y*) normalized by the initial mass *M*_0_. Inset: Temporal evolution of the spatial variance of fluorescein mass in the *y*-direction, 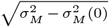, for the 3 cases. The yellow, cyan and purple arrows superimposed on the images depict the width 2*σ*_*M*_ of the Gaussian function that would fit the *y*-profile of the three mass fields.

To understand how food detection by cells occurs at an increasing distance from the injection point, and to reveal any potential flow, we perform a passive solute tracer experiment by injecting a mixture of yeast and fluorescein-dyed water among a group of *S. lemnae*. To do so, a few tens of cells are transferred to an observation chamber consisting of two parallel glass plates separated by a 3 mm gap, large enough to avoid any confinement effects (39). The dyed mixture is injected using a syringe pump in the chamber just above the bottom plate. Imaging the evolving mass field of fluorescein is accomplished using LED-Induced Fluorescence (LEDLIF), where the fluorescence intensity field *I*(*x, y*) in the raw images is converted to the mass field *M* (*x, y*) of fluorescein accumulated in the *z*-column above each pixel (x,y), via the linear relation *M* (*x, y*) ∝ *I*(*x, y*) (see section 4 of *SI Appendix* for the full procedure).

The experiment probing motion of solute tracer among the cells in the presence of food is compared to two other tracer experiments: i) a purely diffusive case consisting of an injection of fluorescein in the observation chamber without cells, and ii) the injection of fluorescein among cells but without yeast. Fig 2C shows images of the dye blobs from each of the three experiments taken 3 min after injection (movie 6). Visible in Fig 2C(ii), the blob injected among cells but without food undergoes some local and intermittent stirring, but its spatial distribution at the end is quite similar to that of the purely diffusive blob seen in Fig 2C(i). By contrast, Fig 2C(iii) reveals how *S. lemnae* cells, in the presence of food, actively generate an outstanding large-scale chaotic flow that, in turn, heavily impacts the shape and mass levels of the blob, transporting fluorescein throughout the observation chamber. The steep mass gradients consistently observed throughout the experiment indicate a Péclet number higher than unity Pe_active_ = *vd*/*D*_fluo_ *>* 1, for a theoretical molecular diffusion coefficient of the fluorescein of *D*_fluo_ ≈ 4.10^−10^ m^2^/s (that is only five times lower than the diffusion coefficient of 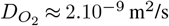). A more quantitative comparison between the tracer experiments can be made by measuring the respective spatial variance of the mass levels in the *y* direction, which is simply the second moment of the mass field: 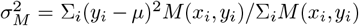, with *µ* = Σ_*i*_*y*_*i*_*M* (*x*_*i*_, *y*_*i*_)*/*Σ_*i*_*M* (*x*_*i*_, *y*_*i*_) the first moment of the mass field and (*x*_*i*_, *y*_*i*_) the position of the pixel. The graphs are shown in the inset of Fig 2B. As expected in the pure diffusive case, the mass variance follows a linear law 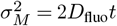, with *D*_fluo_ the theoretical diffusion coefficient expected for fluorescein. Case ii) displays a very similar scaling. Whereas in case iii) where both cells and food are present, after an initial transition time 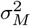 a super-diffusive scaling 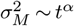 with *α >* 1 for over a decade in time follows until *t* ≈ 200, s, when the blob finally reaches the boundaries of the observation chamber. The ratio of characteristic spreading lengths *σ*_*M*_ between tests i) and iii) reaches a maximum of about ∼ 6 as shown in the figure. This experiment demonstrates how only a few tens of *S. lemnae* can overcome the limits imposed by Stokes conditions to generate a collective flow - triggered by food detection - that spectacularly moves, spreads, and stirs any substances either dissolved or suspended in water. One can extrapolate this observation to natural situations where potentially much larger clusters composed of many more individuals form.

## 3. Hydrodynamic disturbances produced by a single *S. lemnae* either fast crawling or anchored

### A. Experimental measurements

How does a collective flow emerge and how does it relate to the switch in locomotion behavior pointed out in an earlier section? Answering this question requires one to first examine experimentally the hydrodynamic disturbances produced by a single cell. We measure and compare the flow in the cell vicinity for the two main locomotion behaviors described previously, i.e. either fast crawling (*v*_*c*_ ≫ 0) or anchored (*v*_*c*_ ≈ 0) on a no-slip boundary. In the observation chamber we inject some additional water containing microbeads used as passive tracers for visualizing the flow. Since we are ultimately interested in fluid transport and mixing, we use PTV which, unlike Eulerian approaches, provides the essential information about net fluid displacement (see section 5A of *SI Appendix* for a description of the method).

The recorded trajectories of microtracers affected by the motion of three fast crawling cells (see movie 7), with crawling speeds around *v*_*c*_ ≈ 1000 − 1500 *µ*m/s, are gathered and shown in Fig. 3A in a single cell frame (see section 5B of *SI Appendix* for a description of the transformation from the lab frame to the cell frame). The tracers are moved over a persistence time of about *T*_*p*_ = 0.5 s. We define the persistence time *T*_*p*_ as the average time during which tracers move due to the cell’s hydrodynamic perturbation ***v***(***x***) (i.e. above the Brownian motion due to thermal agitation) which for large *v*_*c*_ varies as *T*_*p*_ ∼ 1*/v*_*c*_. Microtracers ahead of the cell are first pumped toward the buccal apparatus before being dragged back in the wake of the cell as it crawls past the tracers. The general picture of the tracer trajectories reveals a very shortrange flow, where the trajectories located on both lateral sides of the cell resemble the closed loops described by (41), in which the displacement of fluid particles by squirmers was modeled. For such trajectories, the net displacement of tracers mostly cancels. A significant net displacement Δ*x* ∼ *L*_*c*_ can essentially be found for the few tracers located on the cell’s path, i.e. for the ones coming into close contact with the stagnation points near the cell body. Indeed, a moving cell can potentially bring in its wake some fluid over even longer distances (‘Darwin drift’), but this is typically limited to a very small volume of fluid (see movie 8). On the other hand, we measure the feeding flow produced by an anchored cell (section 5C of *SI Appendix* and movie 9), an example of which is displayed in Fig. 3B. In this last example, lasting *T*_*p*_ = 7.6 s, the flow is uni-directional, and the net tracer displacement reaches distances up to 5-6*L*_*c*_. We can qualitatively see from the direct comparison between the two cases that the long-range feeding flow is considerably more efficient at transporting fluid over long distances, and this is all the more true as the cell remains motionless and feeds. While feeding, *S. lemnae* remains motionless on average for a few seconds, and may reach pumping times of about ∼100 s.

**Fig. 3.**
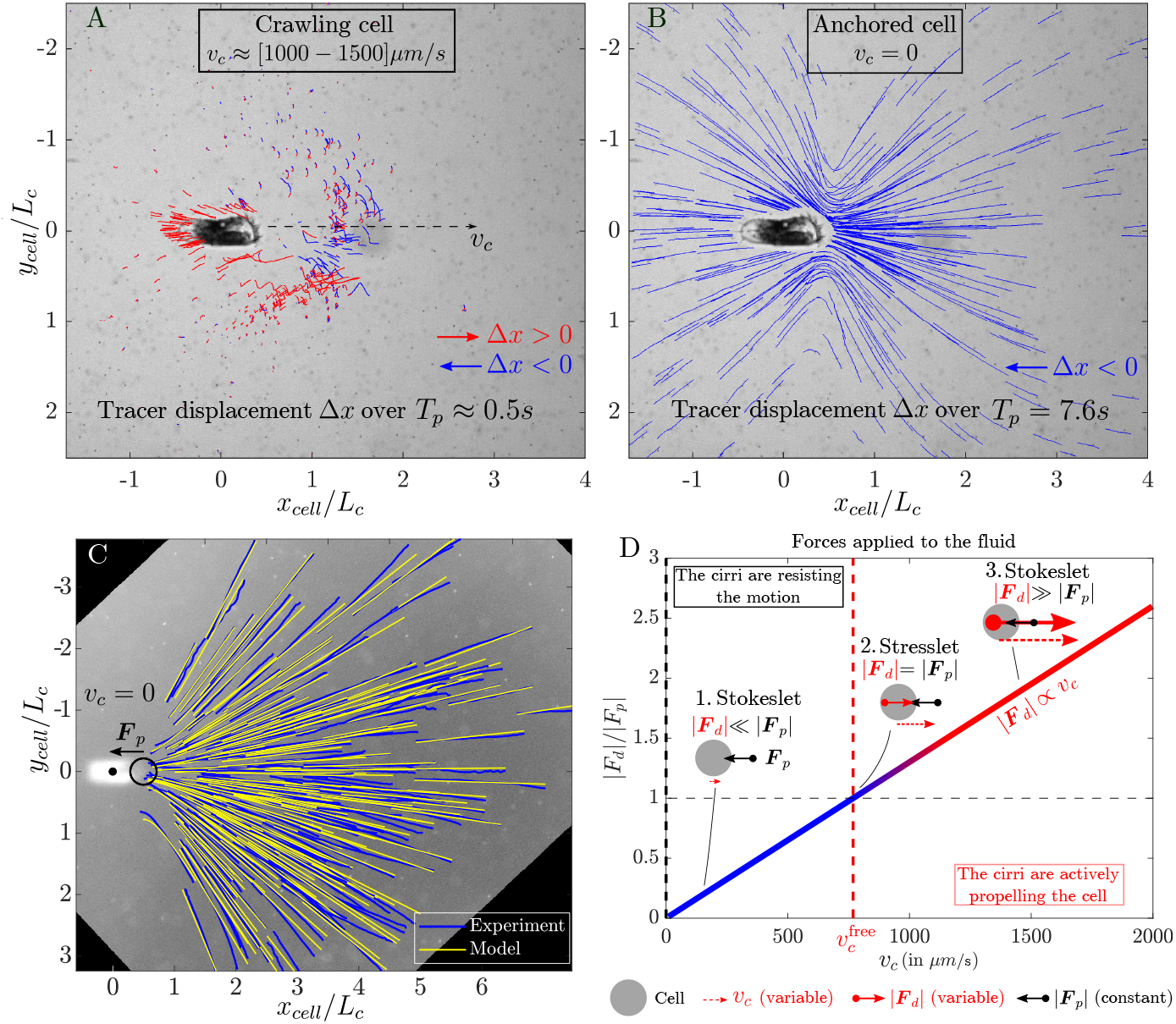
Comparison of the hydrodynamic disturbances caused by an isolated *S. lemnae* either (A) crawling (*v*_*c*_ ≫ 0) or (B) immobile (*v*_*c*_ ≈ 0), projected in the *xy* plane, in the cell frame. In the crawling case (A), fluid particles are displaced over a persistence time about *T*_*p*_ ≈ 0.5 s. In the immobile case (B), the cell remains anchored for *T*_*p*_ = 7.6 s. In both cases, fluid particles trajectories are colored blue if they move left (Δ*x <* 0) or red if they move right (Δ*x >* 0). (C) Comparison between the experimentally measured streamlines (in blue) ahead of a feeding cell and the theoretical streamlines predicted by the model of (40) (in yellow), in the *xy* plane. The black dot marks the center of the ellipsoid shape matching the cell’s body ***x***_*c*_ = (0, 0), and the black circle indicates the position ***x***_*f*_ and radius *δ* of the volume force. (D) Relative importance of the magnitude of the two opposite forces applied to the fluid by the action of the cell (depicted as a gray disk), i.e. drag force *F*_*d*_ vs pumping force *F*_*p*_. *F*_*p*_ is assumed to be constant, while *F*_*d*_ scales with speed as *F*_*d*_ ∝ *v*_*c*_. Depending on the ratio *F*_*d*_*/F*_*p*_, three distinct scenarios for the far-flows are identified: 1) Immobile or slow-moving Stokeslet above a wall (*v* ∼ 1*/r*^2^), with ***F***_*p*_ opposing to direction of cell locomotion. 2) Stresslet above a wall (*v* ∼ 1*/r*^3^), at the terminal velocity 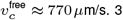) Fast-moving Stokeslet ***F***_*d*_ aligned with the direction of cell locomotion (*v* ∼ 1*/r*^2^).

### B. Modeling the individual feeding flow by a regularized Stokeslet above a no-slip plane

We model the single feeding flow by a regularized (volume) Stokeslet above a wall, from the theory developed in (40, 42) (*SI Appendix*, section 5D). The model needs five input parameters: the location of the force ***x***_*f*_ in the *xy* plane, its height *h* above the wall, the radius of the force *δ*, the orientation of the force in 3D ***F***_*p*_ / |***F***_*p*_| = (cos *θ* cos *ϕ*, sin *θ* cos *ϕ*, sin *ϕ*) (where *θ* and *ϕ* are, respectively, the angle with respect to the *x* axis in the *xy* plane and the angle with respect to the wall *z* = 0) and its magnitude *F*_*p*_ = |***F***_*p*_|. The parameters ***x***_*f*_, *h, δ* and *ϕ* are fixed and extracted from the anatomical features of *S. lemnae* (see the scheme of Fig. 1B). The force position is set to 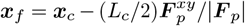 with ***x***_*c*_, *L*_*c*_ and 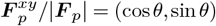 the center of the cell, the average length, and the force orientation in the *xy* plane, respectively. The parameter *h* is the height of the cell’s anterior above the plane and the force radius *δ* is taken as the halfwidth of the cell. The angle *ϕ* with respect to the *z* = 0 wall is measured from the systematic inclination of the cell’s anterior toward the top. In summary, we measure on average *L*_*c*_ ≈ 200 *µ*m, *h* ≈ 50 *µ*m, *δ* ≈ 43 *µ*m, and *ϕ* ≈ 13^°^. These values will remain constant for all cells, ignoring the slight variation in their size. Only the cell position ***x***_*c*_ and orientation *θ* vary and will be extracted systematically from the images. In Fig. 3C, analytical streamlines are compared to a set of experimental streamlines consisting in nine individual feeding flows superimposed all together, produced by nine different cells varying slightly in size, with *Lc* = 200 ± 20 *µ*m (see section 5C-E of *SI Appendix* for the complete validation procedure). Visual comparison between analytical and experimental streamlines shows a nearly perfect geometrical match, thus validating the model. Note that the flow is inherently 3D, as the boundary wall imparts strong parallel shear and bending of the streamlines in the transverse *z* component (see Fig. S3A in *SI Appendix*). Last, the force magnitude |***F***_*p*_| is determined by finding the best match between the magnitudes of the experimental and analytical velocity fields using the least squares fitting. We get |***F***_*p*_| = 1200 ± 240pN. This value seems consistent compared to those of other attached filter feeders found in the literature and is plotted as a function of cell dimension, see Fig. S4 of *SI Appendix*. Note that the residual error on |***F***_*p*_| is mainly due to the finite-size distribution of cell lengths *L*_*c*_, and we expect it to be the main source of error for computing the total collective flow as well.

These velocity field measurements allow us to provide estimates for both Reynolds and Péclet numbers. The Reynolds number is *Re* = *vL/ν*, with *L* the characteristic feeding flow size, *v* the fluid velocity and *ν* ≈ 10^−6^ m^2^/s the kinematic viscosity of water. Taking *L* ∼ 500 × 10^−6^ m and *v* ∼ 100 × 10^−6^ m/s, we get *Re* = 0.05 ≪ 1, ensuring Stokes conditions. The Péclet number related to the fluorescein experiment shown in Fig. 2 is estimated to be Pe_active_ = *vL/D*_fluo_ ∼ 125. Regarding oxygen, we have Pe_active_ ∼ 25. Given the wide range of molecular weights of dissolved organic matter in aquatic environments with diffusivity ranging from *D* = 10^−12^ to 10^−9^ m^2^/s (43), the associated Péclet number ranges from Pe_active_ = 50 up to 50000, giving a first hint of the importance of feeding flow over diffusion for many types of nutrients.

### C. The locomotion speed determines the type of hydrodynamic disturbance

We explain the origin of the two hydrodynamic disturbances using the balance of forces applied to the cell while crawling at a given speed *v*_*c*_ (see the scheme in Fig. 1B). The drag force ***F***_*d*_ opposes two propulsive forces, i.e. the pumping force ***F***_*p*_ generated by its AZM and the contact force ***F***_*c*_ between the cirri and the solid substrate. Hence, for a cell crawling on a straight line at constant speed, we have the balance of forces ***F***_*d*_ = − (***F***_*c*_ + ***F***_*p*_). Since the drag force is proportional to the locomotion velocity ***F***_*d*_ = − *C*_*d*_***v***_***c***_ (with *C*_*d*_ the drag coefficient whose value is estimated in section 6 of *SI Appendix*), the locomotion velocity ***v***_***c***_ can be deduced from the last equation:

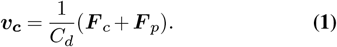

In which proportion do these two forces contribute to *v*_*c*_? Based on observations that the cell AZM appears to be consistently in action during both crawling and pumping events, we assume that the magnitude of the pumping force |***F***_*p*_| = 1200, pN remains constant. Only during abrupt SSR events or mitosis does the AZM seem to stop. In the event that ***F***_*p*_ was the only propulsive force and ***F***_*c*_ = 0, ***F***_*d*_ would balance ***F***_*p*_ at the terminal velocity *v*_*c*_ ≈ 770 *µ*m/s (see again section 6 of *SI Appendix*). Yet, *S. lemnae* can achieve much greater locomotion speed, up to *v*_*c*_ ∼ 7000 *µ*m/s, as can be seen on Fig. 1F, which suggests that the force ***F***_***c***_ plays a driving role in the crawling mode.

How do these forces impact the surrounding fluid? While three forces act on the cell, only two forces are applied by the cell to the fluid, i.e. the opposite forces ***F***_*p*_ and ***F***_*d*_. In fact, the contact force ***F***_*c*_ between the cirri and the wall is assumed here to be transmitted to the substrate in full and to have no effect on the fluid, so that the far field flow is set by the relation ***F***_*d*_ = −***F***_*p*_. However, *F*_*c*_ = |***F***_*c*_| as an external force ultimately determines the amount of drag force *F*_*d*_ = |***F***_*d*_|, since *F*_*d*_ ∝ *v*_*c*_ and according to Eq. (1) *v*_*c*_ ∝ *F*_*c*_. As a consequence, we end up with three different possible scenarios of far-field flows produced by the cell, depending on the relative importance of *F*_*d*_ over the constant *F*_*p*_, as summarized in Fig. 3D. When the cell is immobile (*v*_*c*_ = 0, see Fig. 3B) or crawling at small speeds such that *F*_*d*_ ≪ *F*_*p*_, it produces an unaltered pumping flow, approximated as a Stokeslet above a wall, decaying as *v* ∼1*/r*^2^ (and opposite to the slow cell motion). When *F*_*d*_*/F*_*p*_ = 1 (that is, about *v*^free^ 770 *µ*m/s), and neglecting the slight inclination of ***F***_*p*_ with respect to ***F***_*d*_, as shown in Fig. 1B, the far field corresponds to a Stresslet (dipole force) above a wall that decays as *v* ∼ 1*/r*^3^ (44), producing fluid trajectories close to those shown in Fig. 3A. In this specific case, no net force is applied on the fluid. Finally, when the cell reaches high speed for which *F*_*d*_*/F*_*p*_ ≫ 1, the far field is again similar to a Stokeslet above a wall, but aligned in the same direction as the cell motion.

### D. Either the cell moves or it moves the fluid

The origin of collective flow lies in the ability of a single hydrodynamic disturbance to displace fluid in ranges Δ*x* comparable to separation distances between feeding cells *d*_*c*_. In fact, the far-field flows pointed out in the previous section induce very different net fluid displacements. The net tracer displacement Δ*x* = |***x*** − ***x***_**0**_| is computed by integrating the advection equation 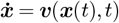 from an initial position ***x***_**0**_:

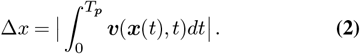

According to the last equation, Δ*x* is influenced by two main parameters: the nature of the vector velocity field ***v***(***x***) (e.g. being unidirectional or not) and its associated spatial correlation (*v* ∼ 1*/r*^2^ and *v* ∼ 1*/r*^3^ for a Stokeslet and a Stresslet above a wall, respectively), and its persistence time *T*_*p*_. Stokeslets are more efficient than Stresslets in fluid transport, as already clear from the comparison between Fig. 3A and Fig. 3B. Then, among the two main scenarios involving Stokeslets (1 and 3 on Fig. 3D), and at constant force magnitude, the persistence time *T*_*p*_ makes the difference: the cumulative net fluid displacement over all fluid particles is the same (the same total volume of fluid is moved) but the average net fluid displacement per particle is not. As a cell moves faster, more particles are affected along its path, but each particle is displaced less (*SI Appendix*, section 7). In brief, the feeding flow range is maximum for *v*_*c*_ = 0, and any increase in *v*_*c*_ reduces it, both because *F*_*d*_ ∝ *v*_*c*_ and *T*_*p*_ ∝ 1*/v*_*c*_ (at least after a transient *v*_*c*_). Thus, the different hydrodynamics observed between Fig. 2C (ii) and Fig. 2C (iii) of section 2 is explained by the average reduction of *v*_*c*_ triggered by food injection, as observed in earlier section. To simplify, either the cell moves or it moves the fluid. Then, considering that the feeding cells remain immobile over *τ*_*c*_ ≈ 2 s on average (as shown later in section 5), in such a case the feeding flow extends up to Δ*x* ∼ 2.5*L*_*c*_ in the *xy* plane. This is comparable to the average separation distance between feeding cells measured in the experiment illustrated in Fig. 2A(ii), which gave ⟨*d*_*c*_⟩ ∼ 4.5*L*_*c*_, so that Δ*x* ∼ ⟨*d*_*c*_⟩*/*2, implying that pumping flows interact hydrodynamically (*SI Appendix*, section 8 for the method to measure *d*_*c*_). In the following collective flow model, we disregard the hydrodynamic contributions of the last 2 scenarios by fixing a maximum locomotion speed threshold of 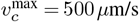. Cells exceeding this speed are considered to exert no force on the fluid.

## 4. Characterization of the collective feeding flow

### A. Analytical skin friction field on the solid boundary

To quantify the strong mixing and dispersion properties of the collective flow observed in Fig. 2, in theory the knowledge about the entire evolving 3D velocity field is required. We here overcome this experimental challenge by demonstrating that the flow can be entirely inferred from the knowledge of its salient topological features, which stem from critical (stagnation) points of the skin friction field (wall shear) on the solid substrate. To illustrate this concept, we consider a simple example where two cells pump in opposite directions, and predict how the resulting flow organizes the spatial distribution of a cloud of tracers (2*µ*m microbeads mixed with *Chlorogonium*-free medium) injected nearby (see Fig. 4A and movie 10). Given the linearity of the Stokes equation, the total velocity field ***v*** at time of image *t*_*j*_ and at position ***x*** = (*x, y, z*) is simply the superposition of all *N* single pumping forces ***v***_*i*_(***x***, *t*_*j*_), depending on varying cell position and orientation (provided 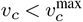):

**Fig. 4.**
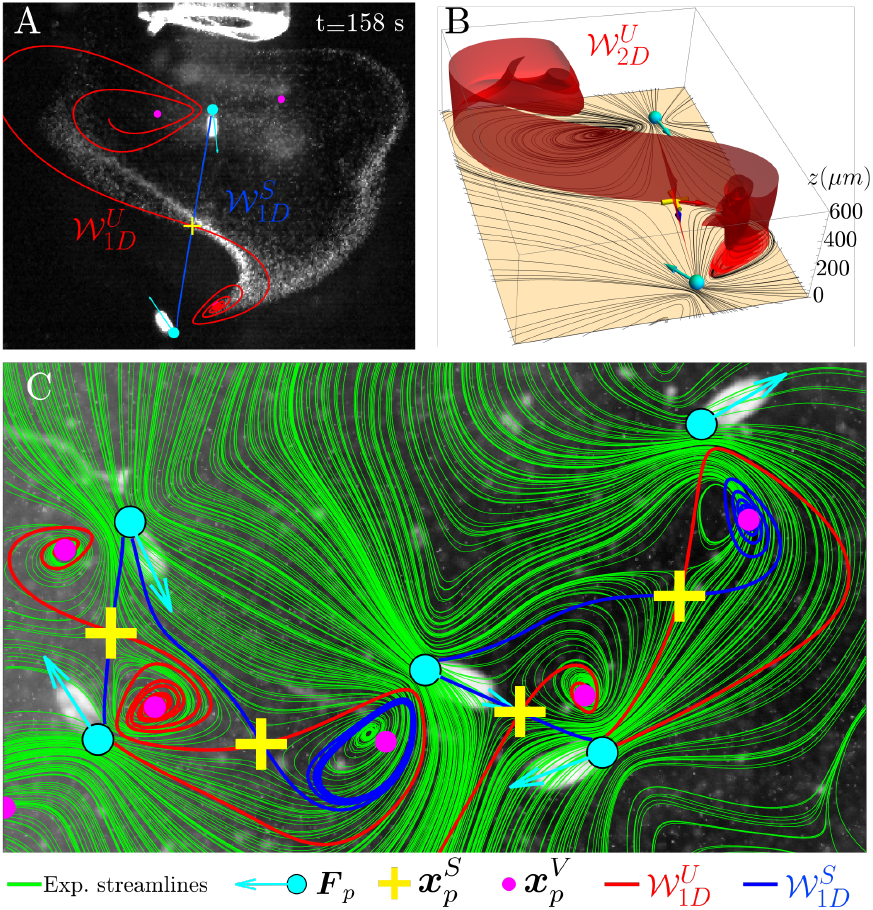
(A) Image of two *S. lemnae* oriented in opposite directions while feeding on a mixture of *Chlorogonium*-free medium and beads (appearing as a cloud of white dots, injected 158 s earlier from the syringe tip visible at the top). Superimposed on the image are the numerical 1D manifolds *W*_1_^*D*^ emerging from the hyperbolic point 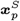 of the analytical skin friction field, computed from the addition of the two volume forces (positions and orientations extracted from the cell images and depicted as cyan dots and arrows respectively). (B) The associated 2D manifold *W*_2*D*_ repelled from *W*_1*D*_ of the skin friction field (whose streamlines are displayed as continuous black lines lying on the *z* = 0 plane) generated by the configuration of cells shown in (A). (C) Instantaneous streamlines (in green) of the velocity field measured experimentally in the *z* = 150 *µ*m plane, produced by a a group of feeding cells. Superimposed on these streamlines are the 1D manifolds *W*_1*D*_ from the associated analytical skin friction field. For clarity, *W*_1*D*_ are truncated whenever they intersect with a volume force.

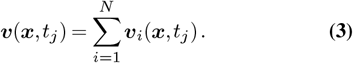

Similarly, since the skin friction field ***τ***_*i*_(*x, y*) produced by a single cell on the solid substrate *z* = 0 is defined as ***τ***_*i*_(*x, y*) = ∂***v***_*i*_/∂*z*|_*z*=0_, the total skin friction field ***τ*** (*x, y*) is again the sum of the skin friction fields produced by every pumping cell 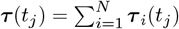. In the example of Fig. 4A a stagnation point must lie on the solid substrate between the two cells, meaning that the skin friction field vanishes at a critical point ***x***_*p*_, defined as ***τ***(***x***_*p*_) = 0. This critical point between the two pumping cells of Fig. 4A is found to be a saddle point 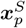 (*SI Appendix*, section 9B). Surana et al. (45) demonstrates that saddle points of the skin friction field are the starting/ending points of 2D manifolds *W*_2*D*_, which act as separation/reattachment material surfaces that propagate in the fluid interior. More precisely, *W*_2*D*_ emanate from critical skin friction lines *W*_1*D*_ connecting two critical points, in our case saddle-spiral connections (see again example of Fig. 4A). The 2D stable/unstable manifolds can be computed numerically by putting non-diffusive fluid tracer particles a small distance above the detected separation/reattachment critical skin friction lines, and by advecting them backwards/forwards in time (see example of *W*_2*D*_ of Fig. 4B, coming from 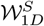 of Fig. 4A). Separation/reattachment surfaces are essential in our system for two reasons: first they provide a set of surfaces of locally minimal flux, i.e. a skeleton for the flow (46) that governs transport; second, because they are also surfaces imposing exponential stretching or compression of nearby fluid elements, they coincide with part of the set of Lagrangian Coherent Structures (LCSs) controlling fluid deformation, and ultimately fluid mixing (47). Getting back to our example seen in Fig. 4A-B, the influence of the 2D unstable manifold 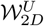 on the spatial distribution of the cloud of tracers can be clearly seen: 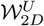 progressively attracts the cloud to the point that it globally fits the shape of 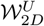, perfectly on the bottom right part of the image, whereas the presence of both the syringe tip and an additional cell just outside the image alters the tracers distribution in the top left corner. Nevertheless, this is an indication that the fate of tracers in the 3D space can be predicted from the knowledge of the analytical critical skin friction lines only. Next we check that the experimental flow patterns can be systematically predicted for the many diverse configurations of cells number, relative positions, and orientations encountered through the imaging. It should be noted that, because our flow is said ‘open’ (the flow intensity vanishes in the *z* direction), all the criteria defined by (45) for reattachment/separation surfaces are not always strictly fulfilled. Anyway, we show in section 9C of the *SI Appendix* that this has little importance as long as we do not look too far from the bottom wall *z* = 0.

### B. Velocity field measurement and comparison with the analytical skin friction field

For each image, the collective velocity field within a feeding cell group is measured using Particle Image Velocimetry (PIV) in a focal plane set at *z*_fp_ = 3*h* above the bottom plate. We inject a mixture of water seeded with microbeads and containing just enough *Chlorogonium* cells to spark the feeding behavior in *S. lemnae*. We get the instantaneous flow by computing the streamlines the curves tangent to the velocity vector field - at each time step (*SI Appendix*, section 9A). An example can be seen in Fig. 4C (other examples shown in Fig. S7 in *SI Appendix*, and movies 11-12). Rich flow patterns can be observed, with a mingling of saddle points and vortices being created, bifurcating and breaking over time, as the cells crawl and reorient. We superimpose on these experimental streamlines the critical points and skin friction lines calculated from the same configuration of cell positions and orientations. Although we compare two sets of streamlines that are at an *z* = 3*h* = 150 *µ*m altitude difference from each other, we assume that the gap distance is small enough so that the sketch drawn by the critical skin friction lines on the wall is representative of the flow patterns in the measurement plane. Within the spatial resolution limits of the velocity field, and provided that no cell outside the image is present to alter the flow inside it, the matching of the two flow patterns is striking, regardless of the cell number, relative separation distances, or orientations. Most likely, the small mismatch that sometimes occurs between analytical critical lines *W*_1*D*_ and experimental streamlines comes from the finite-size distribution of cell sizes, not included in our model. We emphasize that our collective flow model is all the more reliable because throughout the experiments, the feeding force never fluctuated, even when the cell reached satiety.

## 5. Dispersion and mixing properties of the collective feeding flow

### A. Cells dispersion vs. fluid dispersion

To further explore the close relationship between locomotion speed *v*_*c*_ and fluid transport efficiency, we quantify the dispersion dynamics of both cells and simulated fluid particles, from the formation of a feeding cluster to its dismantling. As shown in Fig. 2A-B, cells initially wandering around soon form a feeding cluster around the yeast injection point. Eight movies lasting 3.5 min each are grabbed approximately every 15 min after food injection at time *t*_0_, up to *t* − *t*_0_ = 109 min. We postulate that the cells’ behavior is stationary over the 3.5 min duration of each movie. The MSDs of cells related to the eight movies are plotted as a function of the internal time of each movie 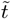 in Fig. 5C. Regarding the first movie *t* − *t*_0_ = 3 min, the MSD tends towards a plateau, in contrast to the Fickian behavior 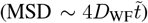 adopted by the cells when in the presence of a uniform food field. This is reminiscent of a chemotactic behavior, where cells manage to stay around the injection point after its detection, in contrast to the random drift effected by cells in a homogeneous chemical landscape, seen in Section 1. At successive times *t* − *t*_0_ we observe a monotonic growth of the MSDs, which ultimately tend to the faster Fickian regime 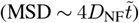, as the accessible food probably becomes scarcer.

**Fig. 5.**
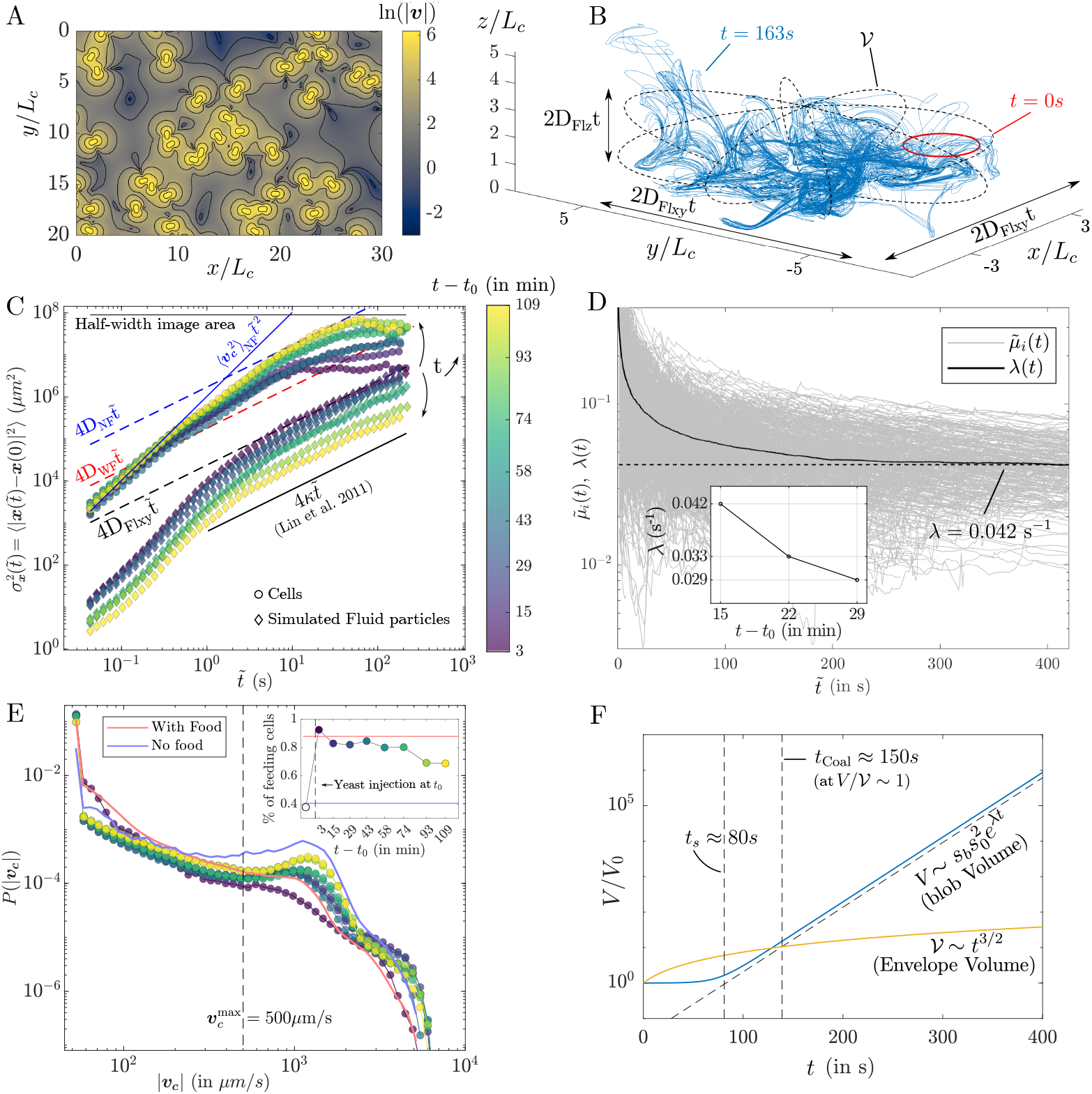
(A) 2D slice of the 3D analytical velocity field at the plane *z* = *h*, computed from the superposition of volume forces over the boundary *z* = 0, with position and orientation extracted from experimental imaging of a feeding cluster at a given time. (B) Material points (tracers) initially arranged on a circle (in red) of length *l*_0_ are advected by the collective flow. After *t* = 165 s the circle is stretched exponentially up to *ρ* = *l/l*_0_ ≈ 650 into the blue material line. (C) Temporal evolution of the MSDs 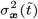 for both cells and tracers (initially arranged on a grid within the cluster). Each curve is computed over 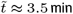, at multiples of (*t* − *t*_0_) ≈ 15 min, with *t*_0_ the injection time. They are compared to the asymptotic MSDs (dashed lines) extracted from Fig. 1H for cells evolving in homogeneous environments (either deprived or immersed in food), as well as to the model of dispersion by squirmers (41), depicted as a continuous black line. (D) Temporal evolution of 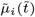, and their average 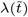 along travel time 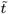, converging to the value *λ* ≈ 0.042 s^−1^, for the movie at *t* − *t*_0_ = 15 min. Inset: *λ* values for the three successive movies at early times *t* − *t*_0_ = [15, 22, 29] min. (E) *P* (|***v***_***c***_|) for successive times (*t* − *t*_0_) after local yeast injection (same colorbar as in (C)), and compared to the homogeneous cases from Fig. 1F. Inset: Proportion of feeding cells over the total number of cells, computed as the fraction of cells with a locomotion velocity 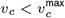. (F) Comparison of the analytic volumes of the blob (in blue, computed with *λ* = 0.042) and the dispersion envelope (in orange). The two volumes intersect at the coalescence time.

To compute fluid dispersion, 2500 material elements (tracers) are initially arranged on a regular grid that covers most of the cluster area (*SI Appendix*, section 10A, and movie 13) set at *z*_0_ = 150 *µ*m. The tracers are numerically advected using the collective flow computed from the evolving cell positions and orientations collected from every movie (see an example in Fig. 5A). We compute both the MSD in the *xy* plane and in the *z*-direction. Note that in our system, the average fluid fluctuations decay with distance *z* from the surface (48). As some tracers are gradually repelled from the surface, the resulting MSDs in both the *xy* plane and along *z* will inevitably decrease, eventually showing a subdiffusive regime. Consequently, here we restrict ourselves to comparing the MSDs relative to each other, bearing in mind that absolute values depend on the integration time. In contrast to cell MSDs, tracer MSDs reveal the opposite temporal dynamics: from time *t* − *t*_0_ = 15 min, a diffusive regime is reached with a maximal dispersion coefficient about *D*_*Flxy*_ = 6.10^−9^ m^2^/s (in comparison, at the same time, *D*_*Flz*_ ∼ 8.10^−10^ m^2^/s = *D*_*F lxy*_*/*7.5, see Fig. S8B of the *SI Appendix*), before a steady decline of the dispersion coefficient through the successive movies. These opposite spreading dynamics are consistent with the results of previous sections, with the cell MSDs mainly depending on locomotion speed *v*_*c*_. The increase in cell MSDs comes from the regular increase of 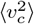, as can be seen directly from the probability distribution of *v*_*c*_ (see Fig. 5E), which shows a regular transfer from the probability of having low *v*_*c*_ towards high *v*_*c*_, above the velocity threshold 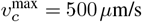 from which in the model the cell does not apply any pumping force. As a result, if the proportion of feeding cells (i.e., for which 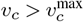) reaches a maximum of approximately 90% just after injection, it then decays to 65%, on the way to the 40% measured prior to injection (see the inset of Fig. 5E). As time goes by, fewer cells contribute to the collective flow, and the tracer MSD decreases. The opposite trends between cell MSD and tracer MSD reflect the remarkable behavioral flexibility of *S. lemnae*, which employs a varying trade-off between chemotaxis and collective flow to swiftly adapt to ephemeral patches of food.

Finally, we compare the simulated fluid dispersion with the model by Lin et al. (41) that gives an estimate of the fluid dispersion coefficient *κ* by a dilute suspension of squirmers moving in an unbounded fluid (*SI Appendix*, section 10B for further details). At equivalent population density (for the movie *t* − *t*_0_ = 15 min) and using the parameters extracted from *S. lemnae* geometry, we find a dispersion coefficient about *κ* ≈ 2.10^−10^ m^2^/s, i.e. approximately 30 times smaller than *D*_*Flxy*_ at *t* − *t*_0_ = 15 min. This 30-fold increase from *κ* to *D*_*F lxy*_ reflects how powerful *S. lemnae*’s collective flow is to transport fluid. Moreover, this value of *κ* is probably overestimated since in our case the no-slip wall accentuates the velocity decay of the Stresslet, from *v* ∼ 1*/r*^2^ to *v* ∼ 1*/r*^3^.

### B. Fluid stretching

While feeding, *S. lemnae* still perform SSRs, implying stochastic changes of orientation. In this way, the 2D manifolds acting as barriers to transport are regularly destroyed while new ones are built. The ensuing unsteady flow imparts stretching and folding of the fluid elements, the hallmark of chaotic advection most favorable to fluid mixing (49), which we quantify here. For example, Fig. 5B illustrates how an initial material line depicted as a red circle is stretched exponentially by the collective flow into a complex lamellar structure (see also movie 14). To quantify the deformation rate, we typically compute the infinite-time Lyapunov exponent, defined as the median stretching rate *λ* = lim_*t*→∞_ ln *ρ*(*t*)*/t* with *ρ*(*t*) = *l*(*t*)*/l*(0) and *l*(*t*) the length of a material line. However, in an open flow the infinite-time Lyapunov exponent is zero. Indeed, at some time all the fluid particles will be expelled along *z* out of the influence zone of the pumping flow, where the velocity fluctuations decay to zero. In other words, the unstable manifold is not space-filling anymore but has a fractal dimension, prior to escaping the influence zone (50, 51), that we call mixing region. Consequently, we rather compute the average of the finite-time Lyapunov exponents (FTLE), evaluated along trajectories as long as they remain inside the mixing region. We define the mixing region as the volume delimited by the minimal envelope surrounding the evolving cluster of feeding cells in the *xy* plane, and by a height limit set to *z*_lim_ (*SI Appendix*, section 11). The choice of *z*_lim_ is delicate as it sets the limit of integration of the FTLE which decays with *z*.

We set *z*_lim_ = 20*h* based on the measurement that only 8% of the fluid particles are lost beyond *z*_lim_ after the 3.5 min duration of the image sequence taken at *t* − *t*_0_ = 15 min, implying that the defined mixing region captures well all Lagrangian kinematics over that period (see Fig. S9B of *SI Appendix*, and also Fig. 5B). Regarding the movie *t* − *t*_0_ = 15 min, we calculate FTLEs 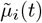 along trajectories *i* within the mixing region (see *SI Appendix*, section 12 for further details about FTLE calculations, and movie 15), and calculate their average 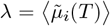. Fig. 5D shows the convergence to *λ* ≈ 0.042 s^−1^. It was possible to obtain the convergence to *λ* for three successive early movies *t* − *t*_0_ = [15, 22, 29] min (note that we incorporate here the movie *t* − *t*_0_ = 22 min, not considered in the previous section), as can be observed on Fig. 5E which shows a decay of *λ* in time, in parallel to the regular decay of the MSDs for the fluid particles. We invoke the same reasoning as in the previous dispersion section to explain the decay of *λ* in time.

Last, the Lyapunov exponent *λ* is made dimensionless as *λτ*_*c*_, where *τ*_*c*_ is the average period during which the cell remains immobile between two successive fast crawling (i.e. 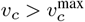) events. Regarding the movie at *t* − *t*_0_ = 15 min, we find *τ*_*c*_ ≈ 2.1 s, which yields a dimensionless Lyapunov exponent *λτ*_*c*_ ≈ 0.09. A dimensionless *λ* allows us to compare the chaotic strength of the flow produced by the cluster of *S. lemnae* to known laminar artificial chaotic flows, and to notice that a collection of feeding *S. lemnae* performs well (for the given mixing region): less than flow through a random pore network (*λ* ≈ 0.117 (52)), random porous media (*λ* ≈ 0.21 (53)), or sheared particulate suspension (*λ* ≈ 0.21 for a solid volume fraction of *ϕ* = 0.35 for example (54)), but better than ordered porous media (*λ* ≈ 0.073 (55)).

### C. Mixing time and coalescence time

We provide here characteristic times indicative of the fluid mixing rate of the collective flow, derived from the lamellar approach (34, 56). This theoretical framework models evolving concentration fields as a set of stretched filaments (lamellae), where the concentration within each lamella is governed by the the coupling between its stretching history *ρ* and molecular diffusion. In 3D flows, mixtures evolve into a set of 3D sheets, whose material area elements align with the principal strain directions, while the compression direction lies mainly perpendicular to the material surface (57, 58). Fluid incompressibility dictates that the sheet width decays as *ρ*^−1^, until enhanced diffusion and compression balance the sheet width down to the Batchelor scale *s*_*b*_, corresponding to the mixing time *t*_*s*_. In chaotic flows, 3D sheets experience a constant elongation rate on average 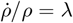 so that *ρ* ∼ *e*^*λt*^. In such case the Batchelor scale remains constant over time as 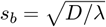, and the corresponding mixing time is *t*_*s*_ = (1*/*2*λ*) ln(1 + 2*Pe*). Of course, in our open flow system, the parameters *s*_*b*_, *λ* and so *t*_*s*_ are in reality space-dependent, varying on average as a function of *z*. Here, we focus on values averaged in the mixing region. Taking *λ* = 0.042, and *D*_yeast_ = 10^−11^ m^2^/s, we get *t*_*s*_ ≈ 80 s. By contrast, the mixing time for a blob of thickness *s*_0_ = 500 *µ*m that is purely diffusing on a still substrate is 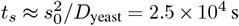, illustrating how powerful exponential stretching is to accelerate dilution. Later, coalescence of the sheets further accelerate homogenization. Initially well separated, the adjacent sheets inevitably merge due to the faster growth of the unfolded blob volume inside a slower-growing dispersion volume (34, 59). Since the material surface grows exponentially while its thickness is locked at the Batchelor scale *s*_*b*_, the unfolded blob volume *V* in turn increases exponentially as:

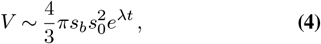

where *s*_0_ is the initial radius of the blob. On the other side, the blob is bounded by the dispersion envelope increasing as a power law:

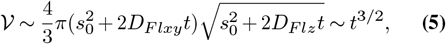

with *D*_*Flxy*_ and *D*_*Flz*_ computed previously. The average number of overlaps is given by the ratio *V/V*. The time at which the lamellae overlap once on average *V/V* = 1 is referred to as the coalescence time, and for the movie *t* − *t*_0_ = 15 min corresponds to *t*_coal_ ≈ 150 s (see the scheme in Fig. 5F). Coalescence further accelerates the homogenization towards the average concentration ⟨*C*⟩ = *C*_0_*/V*. It is important to note that the characteristic mixing and coalescence times occur very early compared to the lifespan of the feeding cluster (∼1 − 2 h in the present case), underscoring the efficiency of the collective flow in mixing solutes. Also, if some fluid is constantly expelled from the mixing region and exchanged with fresh fluid, at *t*_coal_ = 150 s only 6% of the fluid particles have left the mixing region (see Fig. S9B of *SI Appendix*). This suggests that, already at *t* − *t*_0_ = 15 min, most cells within the cluster probe a similar concentration of food or nutrients. Additionally, a cell arriving at the edge of an already established feeding cluster has in a instant a fair appreciation of the chemical content within the mixing region.

## 6. Discussion: a feeding cooperative behavior?

By organizing in a feeding cluster, *S. lemnae* gains the ability to spread and mix fluid on larger scales. One can legitimately ask whether this organization provides benefits to *S. lemnae*, and at what cost: in other words, is the feeding cluster of a cooperative nature? For now, improved foraging success is the notable benefit evident from our results. Indeed, the active spreading of food by the formation of an ever-growing feeding cluster increases the probability that starving cells may locate it, scaling with the patch surface 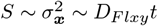. Moreover, we find that food spreading can be even faster when the feeding cluster is already established: we performed the simple experiment of a second yeast injection near a feeding cluster established earlier by a first food injection (*SI Appendix*, section 13A and movie 16). From the second injection time, we computed both cell and tracer MSDs as in section 5 (*SI Appendix*, section 13B). While the cell MSDs display a subdiffusive behavior indicative of some chemotaxis response, the tracer MSDs exhibit a persistent superdiffusive regime over 2.5 decades in time as 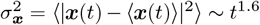. This anomalous scaling of 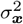 is indicative of persistent hydrodynamic interactions, leading to efficient food transport through preferential flow paths. More generally, established feeding groups can serve as effective substrates for the dissemination of information, such as warning about predators or resources. Other potential physiological benefits can be envisioned. Since many bacteria and protists thrive at narrow interfaces between different chemicals or nutrients at a specific concentration (like at the oxic/anoxic interface (25)), collective fluid spreading could widen the spatial extent of ideal conditions at the population scale. Similarly, it could work at diluting toxic repellent chemicals produced by preys below a given harmful threshold. Furthermore, chaotic mixing quickly smooths out the local depletion areas caused by cell respiration or feeding, thereby sharing resource more efficiently. Finally, by mitigating crowding effects (and so, local competition), chaotic mixing likely boosts the feeding rate per cell, a point that remains to be quantified experimentally.

With regard to the costs of maintaining collective flows, our preliminary estimate is that they are not significant. As mentioned above, collective flows arise primarily from the reduction in locomotion speed *v*_*c*_ of cells clustered around a food patch. Modulating locomotion velocity is a well-known chemotactic strategy that is sufficient to explain the individual behavior of cells. Furthermore, near a food patch, *S. lemnae*’s behavior is remarkably similar whether isolated or in a cluster. In all cases, the cells remain in the food patch, the feeding flows do not weaken, and they persist beyond the point of cell satiety. The most notable distinction arises from the physical contacts that inevitably occur as cells cluster together. These contacts elicit avoidance reactions, which manifest primarily as SSR moves that can propagate throughout the population. Far from having a negative impact, this effect may help maintain an average minimal distance between cells within the cluster, allowing for more efficient space coverage by the cluster. Whether this phenomenon represents a form of self-organization will be the subject of future research.

Thus, since the benefits appear to outweigh the costs, we believe that the feeding cluster represents a form of cooperative behavior employed by *S. lemnae*. Until further investigation potentially uncovers more active forms of cooperation such as active signaling or enhanced feeding rates per cell, we recognize here a type of intra-species cooperation termed byproduct cooperation. This cooperation behavior might extend to other species of Spirotrich, as we have observed similar behavioral patterns in *S. mytilus* and *E. vannus*. More broadly, the hydrodynamic cooperation observed in *S. lemnae* share striking similarities with that of previously cited organisms, i.e. bacteria *T. majus* and *Uronemella* (28–30): in both cases, clusters form in response to a resource gradient, leading to long-range flows. Similarly, just as collective flows in *T. majus* and *Uronemella* depend on variations in population density driven by individual random walks, chaotic mixing in our system would not occur without stochastic reorientations of the cells. The main difference lies in the lack of physical bonds between individuals in the *S. lemnae* cluster, which likely enables them to adapt faster to dynamic environments and efficiently exploit localized ephemeral patches. In contrast, colonies such as *volvox* or choanoflagellates, which take hours to form, would be better suited for larger and persistent patches.

To reconcile predictions of feeding rates with experimental observations for different cell organizations in a patchy food environment, our results suggest that the non-steady nature of the feeder’s motion plays a crucial role, as discussed in the Introduction. Not only does this dynamic stochastic motion affects chemotaxis, but it also allows chaotic flows to occur. In fact, the oscillations observed over time between members of a *Stentor* cluster (60), the erratic contractions of *Vorticella* (8), the alternating cilia beat regime of *O. asymmetrica* (16), or the stochastic and uncorrelated beating of flagella in *S. Rosetta* colonies (61), may just be additional examples of stochastic motions primarily evolved to induce chaotic flow, thus enhancing feeding efficiency. Future studies will benefit from incorporating cell stochastic motion into their models, as ignoring it may impose overly restrictive topological constraints on the Lagrangian kinematics of active flows and preclude chaotic advection.

## 7. Conclusions and perspectives

We described the cooperative foraging behavior of *S. lemnae* in a patchy environment. Upon discovering a food patch, *S. lemnae* reduces its locomotion speed *v*_*c*_, thus increasing the range of its feeding flow. Because cells cluster around the patch, individual feeding flows interact hydrodynamically, leading to a collective flow on the population scale. Despite successfully ingesting food, the cells continue to perform stochastic reorientations, thus imparting chaotic advection. By combining experimental and numerical methods, we measured and predicted the full 3D unsteady flow, and quantified its transport and mixing properties. Our results show that the feeding cluster is at least 30 times more efficient at spreading fluid than a suspension of swimmers, and that the Lyapunov exponent of the cluster’s flow is comparable to that of artificial chaotic flows. Given the substantial benefits granted by collective fluid mixing without additional apparent costs, and the observation that *S. lemnae* behaves similarly whether isolated or in a group, we propose that the feeding group exemplifies a form of intra-species cooperation known as by-product cooperation. This study establishes a quantitative framework for understanding the highly dynamical interplay between an organism’s behavior and the spatial distribution of a resource that it actively shapes. In particular, we hypothesize that optimal feeding for cell clusters in patchy environments relies more on fluid stirring patterns than on simple environmental scanning flux.

Many aspects of a feeding cluster remain to be explored. For instance, how might the cluster adapt its behavior to varying food properties (e.g., food dispersion coefficients), different surface architectures (like porous media) or diverse ambient conditions (such as shear flow, uniform background flow, or confinement)? Additionally, does the cluster organization enhance the average feeding rate per cell? Could it reflect underlying coordinated cooperative behavior, such as quorum sensing, driven by active chemical signaling? How might the cluster respond to the detection of a predator’s repellent? These are just examples of questions we plan to investigate in future research.

## ACKNOWLEDGEMENTS

We are deeply grateful to Dieter Ammermann for providing us with *S. lemnae*. We thank Elena Di Poi for backup cultures. R.T. thanks Nicolas Salles, Nina Javerzat, Benjamin Walter and Bloen Metzger for helpful discussions. We gratefully acknowledge the support by the European Research Council through the ERC PoC Grant Stripe-o-Morph (GA 101069436) and by the Italian Ministry of Research through the projects Response (PRIN 2020), Abyss (PRIN 2022) and Green-Swarm-Intelligence (PRO3).

