## Supplementary Information for "Cooperative mixing through hydrodynamic interactions in *Stylonychia lemnae*"

### 1. Material and methods

**A. Culture of cells.** Strains of *S. lemnae* were provided by Dieter Ammermann (Tübingen, Germany). Cultures were maintained in sterile, HCl-washed glass bowls (Arcoroc, France) containing 10 ml of Eau Volvic a $15°$C at a light:dark cycle of 12:12 h and were fed twice a week with the algae *Chlorogonium sp.* and occasionally with suspended yeast. Fresh cultures were established every two weeks transferring about 30 cells into new culture medium. For experimentation, cells were transferred to a Petri dish and adapted at $20°$ C while maintaining the other conditions stable. Finally cells were starved for a minimum of 24 h prior to every experiment.

**B. Experimental setups.** The imaging is performed directly within the Petri dish culture or inside an observation chamber consisting of two parallel glass plates separated by a $H = 3$,mm gap. This gap is designed to be large enough to prevent the formation of artifact vortices, which otherwise occur on both sides of the pumping cell under confinement. Specifically, such vortices occur when $H/h \sim 1$, where $h$ is the distance from the force singularity to the bottom wall. As Liron theorized (1), the influence of the second wall on the flow can be neglected starting from $H/h > 8$. In our setup, we measure $h = 50\,\mu$m from the average height of the AZM (the location of the singular force), so that $H/h = 60$. To suppress spurious background flow caused by evaporation in the observation chamber, humidity is maintained at saturation thanks to two water tanks placed on either side of the observation chamber, and the entire setup is sealed with a Petri dish cover.

### 2. Sensing food triggers a change of locomotion behavior in *S. lemnae*

**A. Experimental method and image analysis.** In a first step, the positions of starving cells crawling on the bottom of their Petri dish culture are tracked for 5 minutes. The population density is kept low to ensure that encounters between cells are rare during their progression on the bottom plate. In a second step, yeast or algae *chlorogonium sp.* is injected into the culture and the mixture is gently stirred by hand for at least 30 seconds until the food concentration is homogeneous throughout the Petri dish. The second sequence of images is then recorded for 5 minutes. This entire procedure is repeated systematically for seven different cultures of the same *S. lemnae* strain.

Imaging of the cells in these seven different cultures was performed by using a Photron Fastcam Mini UX100 camera mounted on a stereo Olympus SZX16 microscope for the no-food condition, and a Basler acA2000 camera mounted on a Optika SZN-T microscope for the with-food condition. Both microscopes were equipped with a 0.7x optical objective. The images were grabbed at 20 fps for the no-food condition and at 16 fps for the with-food condition.

Cells were detected and tracked using custom Matlab programs, and their trajectory was reconstructed using the Matlab particle tracking velocimetry (PTV) code by Blair and Dufresne (2). In total, we collected about $1 \times 10^5$ and $1.4 \times 10^5$ velocity data points $v_c$ for the conditions without food and with food, respectively. Over a 5-minute period, this corresponds to tracking 330 and 466 cells simultaneously.

**B. Computation of the dispersion coefficient for crawling cells through the Green-Kubo relation.** Because the asymptotic behavior of the mean-square displacement $\sigma_x^2(t)$ is limited by the finite size of the system, we calculate it using the velocity autocorrelation function (VAF). Assuming that the cell locomotion behavior does not vary statistically over the duration of the video, the VAF is stationary. By inserting the displacement equation for a cell $\boldsymbol{x}_c(t) - \boldsymbol{x}_c(0) = \int_0^t \boldsymbol{v}_c(t')dt'$ into the variance of displacement equation $\sigma_x^2(t) = \langle |\boldsymbol{x}_c(t) - \boldsymbol{x}_c(0)|^2 \rangle$ we obtain:

$$\sigma_x^2(t) = \int_0^t dt' \int_0^t dt'' \langle \boldsymbol{v}_c(t') \cdot \boldsymbol{v}_c(t'') \rangle, \quad (1)$$

where $\langle \boldsymbol{v}_c(t') \cdot \boldsymbol{v}_c(t'') \rangle$ is the velocity autocorrelation function, which describes the correlation between the velocities of the cell at two different times $t'$ and $t''$. The angular brackets denote an ensemble average over all times t for a single cell trajectory and over all cell trajectories $i$. Since the system is assumed to be stationary in the long-time limit, the autocorrelation function is independent of the absolute time and depends only on the time lag $\tau = t' - t''$. Therefore, we can express $\langle \boldsymbol{v}_c(t') \cdot \boldsymbol{v}_c(t'') \rangle$

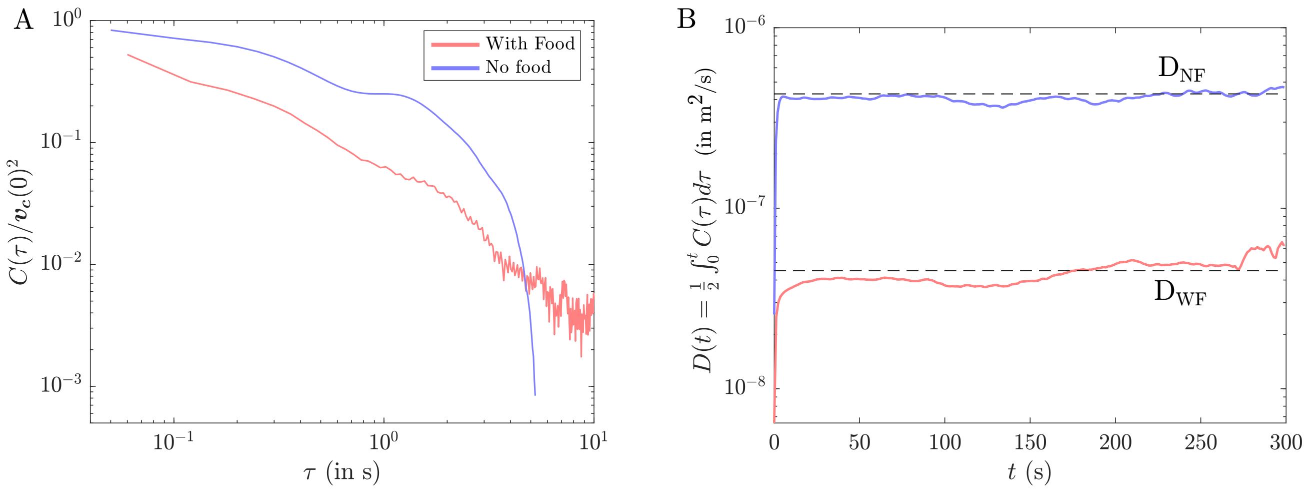


**Fig. 1.** (A) Velocity auto-correlation function $C(\tau)$ as a function of the time lag $\tau$ (Fig. 1F of the main text). (B) Temporal evolution of $D(t)$ computed from the graphs of panel (A) and estimation of the cells dispersion coefficient $D$ (shown as black dashed lines) in both cases.

as $\langle \boldsymbol{v}_c(\tau) \cdot \boldsymbol{v}_c(0) \rangle$. The graph of $C(\tau) = \langle \boldsymbol{v}_c(\tau) \cdot \boldsymbol{v}_c(0) \rangle$ for both cases is shown in Fig. 1A and Fig. 1F of the main text. Using the change of variable $\tau = t' - t''$, we can rewrite Eq. 1 as:

$$\sigma_x^2(t) = 2\int_0^t d\tau (t-\tau)\langle \boldsymbol{v}_c(\tau) \cdot \boldsymbol{v}_c(0) \rangle. \quad (2)$$

The dispersion coefficient $D$ is defined as:

$$D = \lim_{t\to\infty} \frac{\sigma_x^2(t)}{2dt}, \quad (3)$$

where $d$ is the dimension of the system (here $d = 2$). Substituting Eq. 2 into this definition:

$$D = \lim_{t\to\infty} 2\int_0^t d\tau \frac{t-\tau}{2dt}\langle \boldsymbol{v}_c(\tau) \cdot \boldsymbol{v}_c(0) \rangle. \quad (4)$$

Since $\frac{t-\tau}{2dt} \approx \frac{1}{2d}$, as $t \to \infty$ we obtain the Green-Kubo relation:

$$D = \frac{1}{d}\int_0^\infty d\tau \langle \boldsymbol{v}_c(\tau) \cdot \boldsymbol{v}_c(0) \rangle. \quad (5)$$

The graph of $D(t) = \frac{1}{2}\int_0^\infty d\tau C(\tau)$ is displayed Fig. 1B. The dispersion coefficient $D$ is estimated from the long-time plateau value of $D(t)$ in each case. Note that $\sigma_x^2(t)$, and consequently $D$, calculated using the VAF may be slightly underestimated, compared to direct measurement of the MSDs. This is because the Green-Kubo relation gives more statistical weight to the longer trajectories, and therefore to the slower cells, which remain in the image longer. In contrast, faster cells tend to exit the field of view more quickly.

### 3. Formation of a cluster around a food injection site

**A. Experimental setup.** The experiment is performed directly in the Petri dish culture. A small amount of yeast is injected from above using a pipette, where it quickly sinks to the bottom. After injection, the Petri dish cover is immediately placed back to minimize evaporation, which could otherwise induce unwanted background flows. We then grab a sequence of images at 20 fps with a Photron Fastcam Mini UX100 camera mounted on an Olympus SZX16 stereo microscope equipped with a 0.7x optical objective, ensuring the widest possible field of view. The exposure time is set to 20 ms, which allows the cell images to remain sharp even at high locomotion speeds $v_c$.

### 4. Solute transport experiment

**A. Experimental setup and imaging of the fluorescent dye.** To obtain the mass fields $M(x,y)$ of fluorescein, we use the LED-Induced Fluorescence (LEDIF) technique, where the dye excited by incident light emits fluorescence at another wavelength with intensity $I$ proportional to its concentration $C$. Because the flow is actually 3D, fluorescein can freely accumulate in the $z$-column above every pixel $(x,y)$, and therefore the measured fluorescence intensity field $I(x,y)$ is not directly convertible to concentration levels, but instead to the mass of fluorescein accumulated inside each column. Ideally, one aims for the best possible signal-to-noise ratio (SNR). However, in our case, we must balance the SNR with the constraints imposed by the cells' natural sensitivity to the maximum incident light intensity and the maximum fluorescein concentration that the cells can tolerate. The optimal compromise is achieved with the following imaging setup: Two LEDs (KL 1600 LED from Olympus), positioned approximately 20 cm apart on either side of the chamber, provide homogeneous incident light illumination. The incident white light is filtered using a short-pass filter ($\lambda = 500$ nm, FESH0500 from Thorlabs), whereas a long-pass excitation filter ($\lambda = 530$ nm, FGL530S from Thorlabs) is used to block the incident light and collect only the fluorescence signal. Under these conditions, the loss of emitted fluorescence intensity due to photobleaching, after 30 minutes of illumination, is measured to be only about $2\%$. A low readout-noise camera (Hamamatsu ORCA fusion BT) mounted on a stereo Olympus SZX16 microscope, combined with a 0.8x objective, provides a resolution of $8.6\,\mu$m/pixel and a field of view of $14.5\,\text{mm} \times 18.1\,\text{mm}$. We capture 16-bit images at 2 fps, with an exposure time of $0.5$ s. With this setup, imaging of the injected blob of dye at $t = 0$ s yields a SNR $\approx 300$ (and a contrast-to-noise ratio CNR $\approx 277$). To ensure linearity between the intensity and concentration of the emitted light, we check that $C \propto I$ is true below the concentration $C_0 = 40\,\mu$g/ml of injected fluorescein. This is done by injecting solutions of increasing concentrations throughout the observation chamber and measuring their corresponding intensity in the images.

**B. Experimental Methods and imaging.** A few tens of cells from a culture starved for 1.5 days are transferred one by one into the observation chamber using a micro-pipette. Since cells are typically stressed after transfer, it is necessary to wait several hours for them to recover and resume normal behavior, which is indicated by their return to high-speed crawling along long leftward arcs. A small amount of yeast is mixed into a solution of Volvic water and dissolved fluorescein with concentration $C_0 = 40\,\mu$g/ml. This dye concentration $C_0$ is the maximum that cells can tolerate, without visibly affecting their locomotion behavior. Using a syringe pump, we inject $V_0 = 0.25\,\mu$l of this solution just above the bottom plate at a flow rate of $Q = 0.05\,\mu$l/s, and then withdraw the needle. Image analysis is performed using custom Matlab programs, converting intensity images $I(x,y)$ to mass fields normalized by the initial mass as:

$$\frac{M(x,y)}{M_0} = \frac{I(x,y) - I_b}{I_0 - I_b}, \tag{6}$$

where $I_0$ is the intensity measured within the dye blob in the first image at $t = 0$, and $I_b$ is the background intensity measured far from the blob.

### 5. Hydrodynamic disturbances produced by a single *S. lemnae* either fast crawling or anchored

**A. Experimental setup.** The observation chamber and cells are prepared as described in the previous solute transport experiments. We then inject additional water containing microbeads with a diameter of $4.5\,\mu$m into the chamber. This injection is performed after the cells recover normal behavior; otherwise, the beads would have time to sediment. The microbead size is chosen to be small enough to accurately follow the flow streamlines, while still large enough to avoid moving by Brownian motion due to thermal agitation. The chamber is placed on an Olympus IX 81 inverted microscope, equipped with a PLN 4X objective associated to a magnification changer of 1.6X to achieve higher magnification of 6.4X. The focal plane is set at $z_{\text{fp}} = 150\,\mu$m and $z_{\text{fp}} = 200\,\mu$m above the bottom plate for the crawling and feeding cases, respectively. We capture sequences of images using a Photron Fastcam Mini UX100 camera (with frame rates of 125 fps and 50 fps for crawling and anchored/feeding, respectively), until we capture each of these two characteristic behaviors several times.

**B. Fast crawling cell: transformation from lab frame to cell frame.** A crawling cell generates a flow located primarily in its vicinity, as can be observed on Fig. 2A, where small microbeads displacements are visible all along its path. To compare the net displacement of microbeads caused by crawling or pumping cells in a single frame, a frame transformation is applied in the case of crawling cells. Assuming that the crawling behavior is consistent over time (with no change in orientation or velocity), there exists a unique set of microbead trajectories determined solely by the position of the first detectable motion of the bead in the image, relative to the cell body. The "first movement" refers to the displacement of the microbead induced by the crawling cell that exceeds its intrinsic Brownian motion (using a threshold on the minimum microbeads velocity).

The position $p^t$ of the first movement of the microbead at time $t$ is detected, as well as the position and orientation of the cell, which together define the cell frame $R^t_{cell}$ (see Fig. 2B). Next, all subsequent positions $[p^{t+1}, p^{t+2}, ...]$ of this microbead trajectory are transformed into this unique initial frame $R^t_{cell}$, regardless of future cell frames in subsequent images. In this way, we end up with trajectories all translated into a single reference frame (Fig. 2C).

The extraction of the cell's position and orientation in each image is performed using a custom Matlab program that fits an ellipsoid to the cell shape. The position of the cell is defined by the ellipsoid's origin, while the orientation is determined by its major axis.

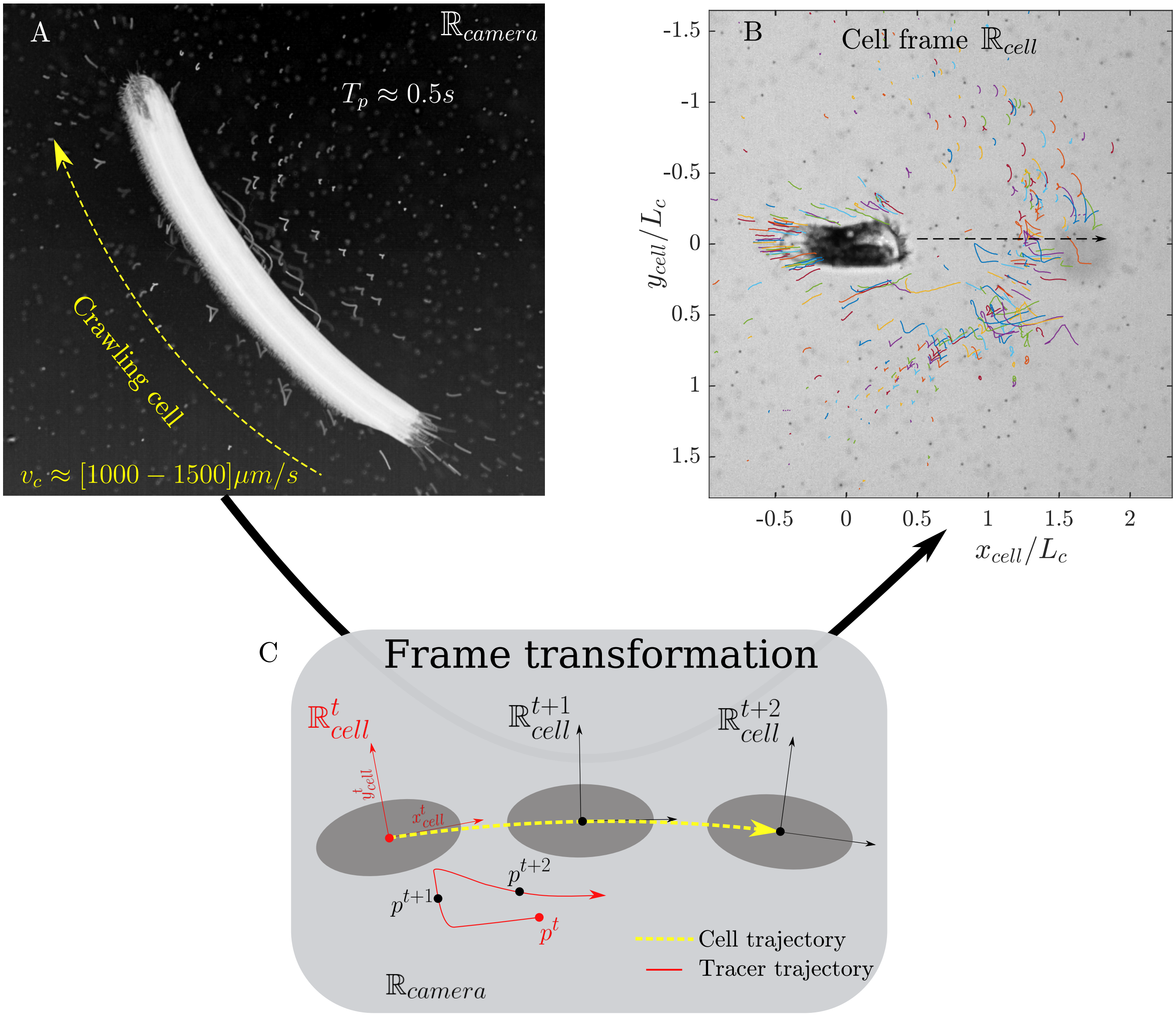


**Fig. 2.** (A) Visualization of the hydrodynamic disturbance in the wake of a cell crawling over $T_p = 0.5$ s using Flowtrace code (3) to compute an image resulting from the stacking of the entire sequence of images. (B) The microbeads trajectories as seen in lab frame (A) are translated to a unique cell frame. (C) Scheme of the frame transformation of the tracers trajectories from lab frame $\mathbb{R}_{\text{camera}}$ (A) to a unique cell frame (B).

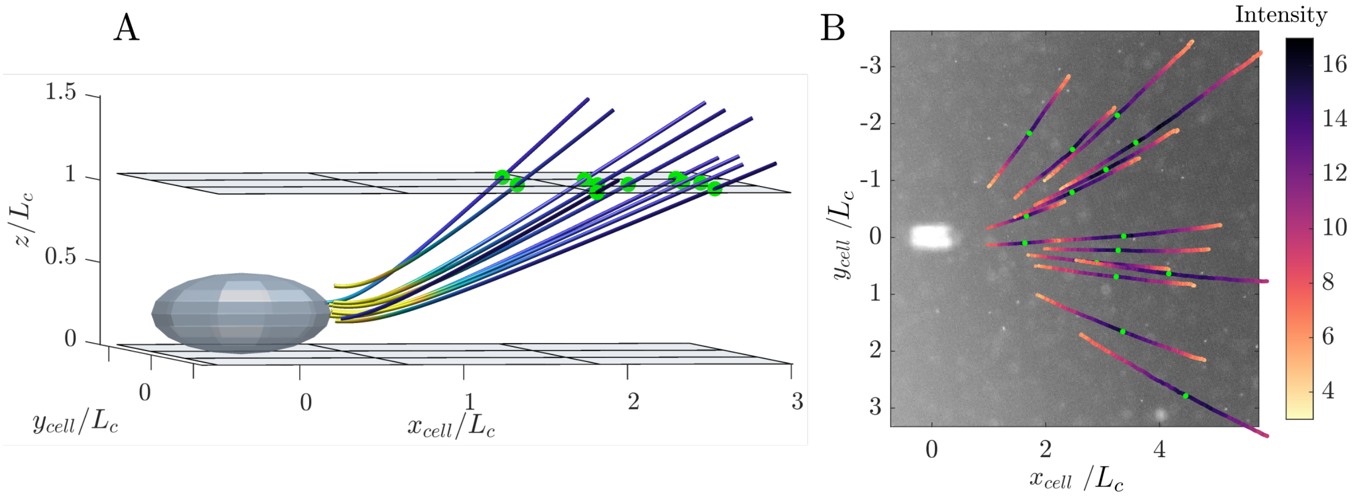


**Fig. 3.** Description of the method used to detect the locations of crossing of the trajectories with the focal plane set at $z_{\rm fp} = 200\,\mu$m, indicated as green dots in both images. (A) A 3D view of some simulated trajectories of the feeding flow produced by an anchored cell represented as a grey ellipsoid positioned at origin $\boldsymbol{x}_c = (0, 0, h)$ (with the volume force ahead of the cell at $\boldsymbol{x}_f = (L_c/2, 0, h)$ ), shows the strong transverse component (in the $z$-direction) of the velocity field. (B) Picture of a few experimental trajectories in the $xy$-plane generated by an anchored cell. The cell is visible in white and centered on the origin $(x, y) = (0, 0)$. Streamlines are colored as a function of the intensity of the successive dots constitutive of the trajectories, directly related to their sharpness is the image.

**C. Hydrodynamic disturbance caused by a single feeding cell: Experimental method.** We measure the streamlines of feeding flows produced by nine individual anchored cells, with a cell length distribution of $L_c = 200 \pm 20\,\mu$m. Each set of experimental streamlines is transformed into a unique cell frame (see blue streamlines in Fig. 3C of main text), where the origin and $x$-axis correspond to the cell center $\boldsymbol{x}_c$ and the cell's longitudinal axis, respectively. In this frame, $\boldsymbol{x}_c = 0$ and $\theta = \pi$ (feeding flow from right to left).

As shown clearly in Fig. 3A, although the feeding force is nearly parallel to the bottom wall, away from the cell there exists a strong transverse component (along the $z$-axis) of the velocity vector. However, the microbead trajectories captured by the camera are projections of their true 3D paths onto the $xy$ plane. To compare the 3D analytical flow with the experimental projected flow, as well as their velocity magnitude, information about the $z$-component of the microbead's positions along the trajectory is required. To address this, we track the evolution of the pixel intensity of the microbeads, as they move with the flow (see Fig. 3B). The intensity of the image microbeads being roughly inverse to their blurriness, the peak of intensity coincides with the point where the microbeads cross the focal plane (when the bead image is the sharpest). These positions are marked with green dots in the figures. Knowing the focal plane altitude (set to $z_{\rm fp} = 200\,\mu$m above the wall), we can then deduce the exact 3D coordinates of the microbeads' positions $\boldsymbol{x}_{\rm fp} = (x_{\rm fp}, y_{\rm fp}, z_{\rm fp})$ at the point where each streamline crosses the focal plane upstream of the cell. Trajectories that do not cross the focal plane are excluded from the analysis. Keeping only the streamlines crossing the focal plane, we are left with 211 microbead trajectories, distributed across the nine different isolated feeding flows.

**D. Modeling the individual feeding flow by a regularized Stokeslet above a no-slip plane.** From the seminal work of Blake (4), the feeding flow of anchored feeding filters is commonly modeled as the flow generated by a force singularity, known as Stokeslet, located at a distance $h$ from a no-slip plane. The velocity field is the solution to the incompressible Stokes equations:

$$\begin{cases} \mu\nabla^2\boldsymbol{v} - \nabla p + \boldsymbol{F}_p = 0\,, \\ \nabla\cdot\boldsymbol{v} = 0\,, \end{cases} \tag{7}$$

with $\boldsymbol{v}$ the fluid velocity, $\boldsymbol{F}_p$ the pumping force applied to the fluid at position $\boldsymbol{x}_f = (x_f, y_f, h)$, $\mu$ the fluid viscosity, and $p$ the pressure field. The no-slip boundary conditions at the wall imposes $\boldsymbol{v}|_{z=0} = 0$. However, the singular nature of the Stokeslet introduces large numerical errors when quantifying fluid deformation. To address this, we instead use a regularized (volume) Stokeslet above a wall, based on the theory developed by (5, 6). The solution is an extension of the system of images for a singular Stokeslet, where the force is concentrated in a "blob" surrounding the point of application. The expression of the blob controls the spatial distribution of the volume force. To represent the force generated by the AZM, we adopt the algebraic blob from (6):

$$\psi(r) = \frac{15\delta^2}{8\pi(r^2+\delta^2)^{7/2}}\,, \tag{8}$$

with $r$ the radial distance from the center of the regularized Stokeslet and $\delta$ the blob radius (see Fig. 1B of main text). The velocity of the fluid evaluated at point $\boldsymbol{x}$ induced by a volume force $\boldsymbol{F}_p$ at position $\boldsymbol{x}_f = (x_f, y_f, h)$ and its image at $\boldsymbol{x}_{f,im} = (x_f, y_f, -h)$ are given by the following.

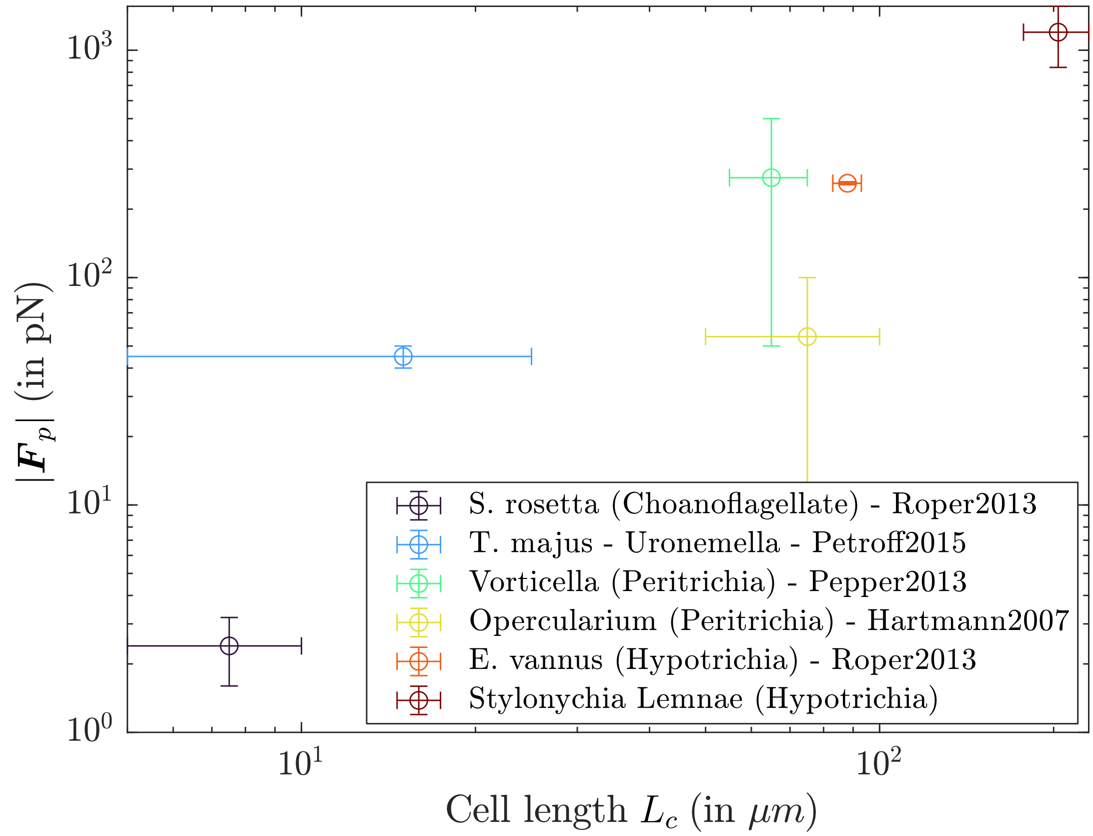


**Fig. 4.** Plot of the pumping force magnitude $|\boldsymbol{F}_p|$ as a function of the cell dimension $L_c$, for different unicellular organisms found in the literature, and comparison with the value measured for our *S. lemnae*.

$$\begin{aligned}\boldsymbol{v}(\boldsymbol{x}) =& [H_1(r^*)\boldsymbol{F}_p + H_2(r^*)(\boldsymbol{F}_p\cdot\boldsymbol{x}_r^*)\boldsymbol{x}_r^*] - [H_1(r)\boldsymbol{F}_p + H_2(r)(\boldsymbol{F}_p\cdot\boldsymbol{x}_r)\boldsymbol{x}_r] \\ & - h^2[D_1(r)\boldsymbol{g} + D_2(r)(\boldsymbol{g}\cdot\boldsymbol{x}_r)\boldsymbol{x}_r] - 2h\left[\frac{H_1'(r)}{r} + H_2(r)\right](\boldsymbol{L}\times\boldsymbol{x}_r) \\ & + 2h\left[H_2(r)(\boldsymbol{g}\cdot\boldsymbol{e}_3)\boldsymbol{x}_r + H_2(r)(\boldsymbol{x}_r\cdot\boldsymbol{e}_3)\boldsymbol{g} + \frac{H_1'(r)}{r}(\boldsymbol{g}\cdot\boldsymbol{x}_r)\boldsymbol{e}_3 + \frac{H_2'(r)}{r}(\boldsymbol{x}_r\cdot\boldsymbol{e}_3)(\boldsymbol{g}\cdot\boldsymbol{x}_r)\boldsymbol{x}_r\right],\end{aligned} \tag{9}$$

where $\boldsymbol{x}_r^* = \boldsymbol{x} - \boldsymbol{x}_f$, $\boldsymbol{x}_r = \boldsymbol{x} - \boldsymbol{x}_{f,im}$, $r^* = |\boldsymbol{x}_r^*|$, $r = |\boldsymbol{x}_r|$, $\boldsymbol{g} = 2(\boldsymbol{F}_p\cdot\boldsymbol{e}_3)\boldsymbol{e}_3 - \boldsymbol{F}_p$ are the dipole strengths, $\boldsymbol{L} = \boldsymbol{F}_p\times\boldsymbol{e}_3$ are the rotlet strengths and

$$H_1(r) = \frac{1}{(8\pi(r^2+\delta^2)^{1/2}} + \frac{\delta^2}{8\pi(r^2+\delta^2)^{3/2}}, \quad H_2(r) = \frac{1}{8\pi(r^2+\delta^2)^{3/2}},$$
$$D_1(r) = \frac{1}{4\pi(r^2+\delta^2)^{3/2}} - \frac{3\delta^2}{4\pi(r^2+\delta^2)^{5/2}}, \quad D_2(r) = -\frac{3}{4\pi(r^2+\delta^2)^{5/2}}.$$

Among the five input parameters required to model the pumping flow produced by an immobile cell (as described in the main text), the only two remaining unknowns are the position $\boldsymbol{x}_f = \boldsymbol{x}_c - (L_c/2)\boldsymbol{F}_p^{xy}/|\boldsymbol{F}_p|$ and orientation $\boldsymbol{F}_p^{xy}/|\boldsymbol{F}_p| = (\cos\theta, \sin\theta)$ of the regularized force in the $xy$ plane. As stated above, in the unique frame, the nine feeding cells share the same position $\boldsymbol{x}_c = 0$ and orientation $\theta = \pi$, so that $\boldsymbol{x}_f = L_c/2$ and $\boldsymbol{F}_p^{xy}/|\boldsymbol{F}_p| = (-1, 0)$.
Next, the analytical streamlines are obtained by solving the advection equation $\dot{\boldsymbol{x}} = \boldsymbol{v}(\boldsymbol{x})$ using the ode45 function in Matlab. We integrate from each unique point $\boldsymbol{x}_{\text{fp}} = (x_{\text{fp}}, y_{\text{fp}}, z_{\text{fp}})_i$ along the streamline $i$ (shown as green dots in Fig. 3A-B), both backward and forward in time, until the length of the analytical streamline matches the length of the experimental one. Finally, the analytical streamlines are displayed as the yellow continuous lines in Fig. 3C of the main text.

**E. Estimate of the Force magnitude.** The measured Lagrangian velocities are the projections in the $xy$-plane of the true 3D velocity field. To compare these experimental velocities with the analytical velocities, we also project the theoretical velocities into the $xy$ plane, so that the magnitude of the theoretical velocity is $|\boldsymbol{v}_{\text{th}}| = \sqrt{v_{x\text{th}}^2 + v_{y\text{th}}^2}$. We compute the analytical velocities

at the 211 specific points where the streamlines cross the focal plane (not shown in the figure). The velocity magnitude varies proportionally to the force magnitude $|\boldsymbol{F}_p|$ applied by the cell. To determine the optimal force magnitude, we perform a least-squares fitting, minimizing the function $\Sigma_i(|\boldsymbol{v}_{\text{exp}}|_i - |\boldsymbol{v}_{\text{th}}|_i)^2$ at all points $i$. The optimal pumping force magnitude is found to be $|\boldsymbol{F}_p| = 1200 \pm 240$ pN (with a minimal residual error of about $18\,\mu$m/s).
Furthermore, we can compare this value to the magnitude of drag force experienced by the cell body while swimming freely. The drag force for a moving sphere is given by the Stokes drag equation: $F = 6\pi\mu\delta v_s$, with $v_s$ the swimming speed. In the images, we measure $v_s = 1300\,\mu$m/s. Using $\delta = 43\,\mu$m, we get $F \approx 1050$ pN, close to the pumping force magnitude $F_p = 1200$ pN.

### 6. Estimation of the drag applied on the body of a crawling cell

Three forces act on the cell during its crawling motion. Two of these forces, the pumping force $\boldsymbol{F}_p$ generated by AZM and the contact force $\boldsymbol{F}_c$ that arises from the interaction between the cirri and the solid substrate, are propulsive. The third force, the drag force $\boldsymbol{F}_d$ opposes the cell's movement (see Fig. 1B of the main text). Therefore, for a cell moving at constant speed, we have:

$$\boldsymbol{F}_d = -(\boldsymbol{F}_c + \boldsymbol{F}_p). \tag{10}$$

We assume that the feeding force magnitude is constant and equal to $|\boldsymbol{F}_p| = F_p = 1200$ pN, as determined previously. The contact force $|\boldsymbol{F}_c|$ is unknown. Here, we provide a theoretical estimate for the magnitude of the drag force. To do this, we model our system as a sphere moving in a quiescent fluid parallel to a wall, with the center of the sphere at a distance $h = 50\,\mu$m from the wall. The radius of the sphere $R$ is taken to be equal to the width of the cell $R = \delta = 43\,\mu$m. The general drag equation is:

$$F_d = \frac{1}{2}\rho v_c^2 A C_d, \tag{11}$$

with $\rho$ the density of the fluid, $v_c$ the velocity of the cell relative to the fluid, $A = \pi\delta^2$ the reference area, and $C_d$ the dimensionless drag coefficient which depends on the system's geometry. For a sphere moving in an unbounded fluid at low Reynolds number $Re$, the drag coefficient is $C_d = 24/Re$, where the Reynolds number is defined as $Re = \rho v_c 2\delta/\mu$, and $\mu$ the dynamic viscosity of the fluid. Substituting this drag coefficient into the drag equation (11) yields the Stokes drag equation $F_d = 6\pi\mu\delta v_c$. When the sphere moves parallel to the wall, a correction is needed to account for the additional drag induced by the wall. Since in our case the gap between the cell and the wall is very small ($h - \delta \approx 7\,\mu$m), lubrication theory applies (7), and we have:

$$C_d = \frac{24}{Re}\left(0.9588 - \frac{8}{15}\ln\left(\frac{h-\delta}{\delta}\right)\right). \tag{12}$$

For $h - \delta \approx 7\,\mu$m, the correction factor is approximately 1.93. Inserting this corrected drag coefficient into the drag equation (11), the drag force is about $F_d \approx 12\pi\mu\delta v_c$, which is found to balance the pumping force $F_d = F_p = 1200$ pN ($F_c = 0$) at the terminal velocity $v_c^{\text{free}} \approx 770\,\mu$m/s. When the cell reaches this speed, and neglecting the small angle $\phi = 13°$ of $\boldsymbol{F}_p$ with respect to $-\boldsymbol{F}_d$ (see Fig. 1B of the main text), the far-field hydrodynamic perturbation generated by the cell can be represented by a stresslet (force dipole) which does not impart a net force on the fluid.
Note that in practice, *S. lemnae* has an elongated shape and a flat ventral side, making it more streamlined than the spherical model used here. Hence, the drag coefficient is probably slightly overestimated, and consequently the free-swimming velocity $v_c^{\text{free}}$ is underestimated.

### 7. Persistence time $T_p$

Fig. 5A illustrates how increasing the locomotion speed $v_c$ of a Stokeslet (of constant magnitude, case of scenario 1 mentioned in the main text) moving along a straight line reduces the net displacement experienced by a given fluid particle. In fact, the net displacement is related to the persistence time $T_p$ (see Equation 2 of the main text), which decays with the cell speed as $T_p \sim 1/v_c$ (see Fig. 5B), after a transient speed (which is $v_c \approx 1\,L_c\,\text{s}^{-1} = 200\,\mu$m/s. According to Fig. 5C, $\Delta x \sim 1/v_c$.
In Scenario 3 where $F_d \gg Fp$, the cell moves at high speed $v_c > v_c^{\text{free}}$ and applies a net force that aligns with its locomotion direction, with a force magnitude increasing as $F_d \sim v_c$. In that case, the net displacement $\Delta x$ of a given fluid particle becomes independent of $v_c$ (but the total volume displaced by the Stokeslet will increase linearly with $v_c$, as it will affect more particles).

### 8. Distance between nearest neighbors

The average distance between the nearest neighbors $\langle d_c \rangle \approx 4.5 L_c$ is calculated by measuring the lengths of the straight lines that connect the feeding cells, using Delaunay triangulation. Only feeding cells - those for which $v_c < v_c^{\text{max}}$ - are considered for the triangulation. An example of such a triangulation can be seen in Fig. 6A. The distribution of the lengths $d_c$ normalized by the cell length $L_c = 200\,\mu$m, calculated for the case shown in Fig. 2A(ii) of the main text, is displayed in Fig. 6B.

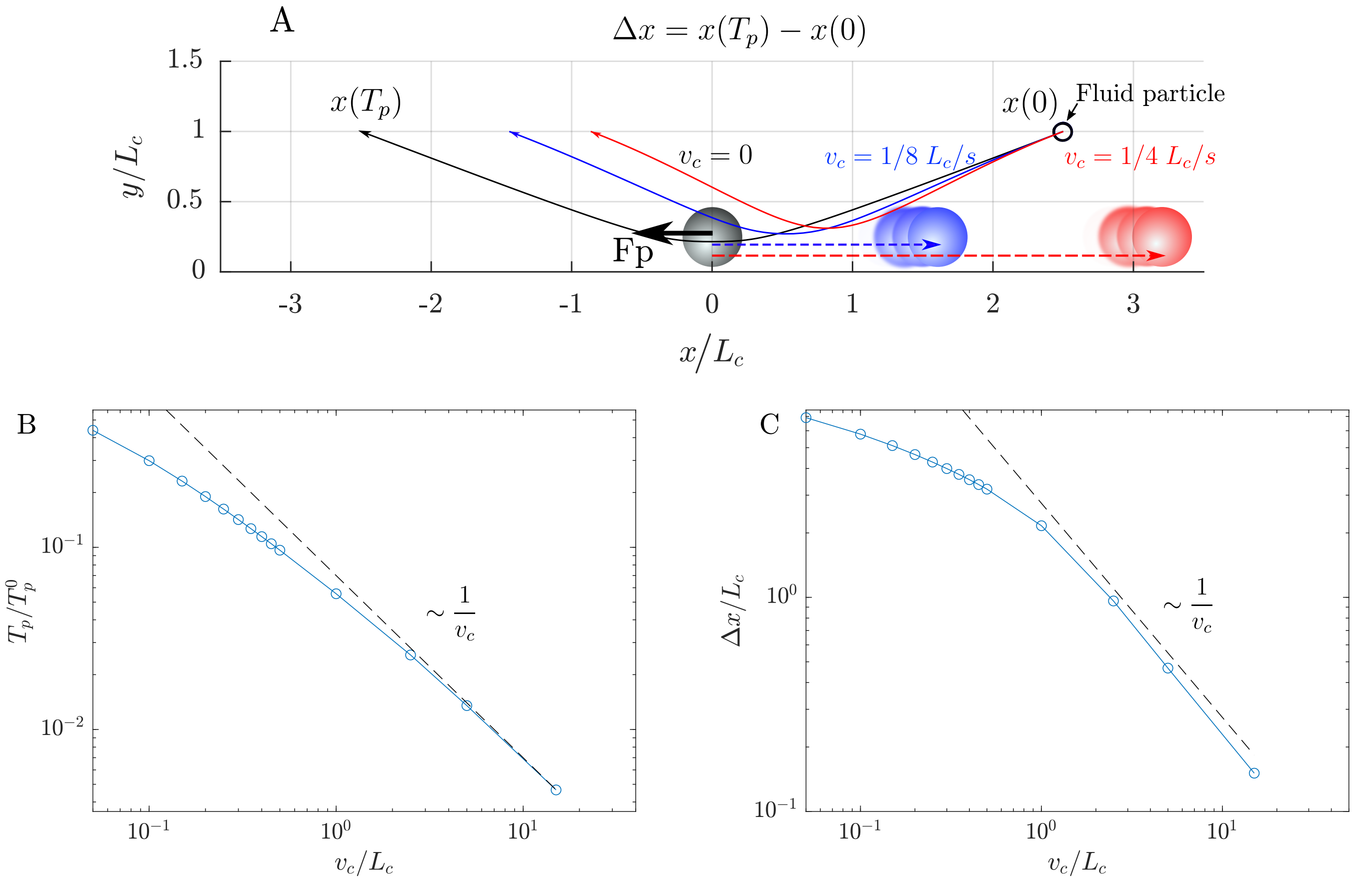


**Fig. 5.** (A) Comparison of several trajectories in the $xz$-plane of one given fluid particle advected by a regularized Stokeslet $\boldsymbol{F}_p$ above the no-slip $z = 0$ boundary and parallel to it ($\phi = 0$) moving on a straight line along the $x$-axis, with three different locomotion speeds $v_c$: either $v_c = 0$ (in black), $v_c = 0.125\,L_c/s$ (in blue) or $v_c = 0.25\,L_c/s$ (in red). The initial position of the tracer is depicted by a small black circle at $\boldsymbol{x}_0 = (2.5, 1)L_c$. All three tracer trajectories are integrated over the perturbation time $T_p$ for which $\boldsymbol{v}(\boldsymbol{x}) > \epsilon$, with $\epsilon = 0.15\,L_c/s$. The initial force position in the $xz$-plane $\boldsymbol{x}_f = (0, h)$ and forcing orientation $\boldsymbol{F}_p$ are depicted as gray disk and black arrow, respectively. (B) Persistence time $T_p$ over which the trajectories are computed, as a function of the cell locomotion speed $v_c$. The initial tracer position is $\boldsymbol{x}(0) = (5, 1)L_c$. $T_p$ is normalized by $T_p^0$, the persistence time for the immobile force case, i.e. $v_c = 0$. (C) Net displacement $\Delta x$ varying with $v_c$.

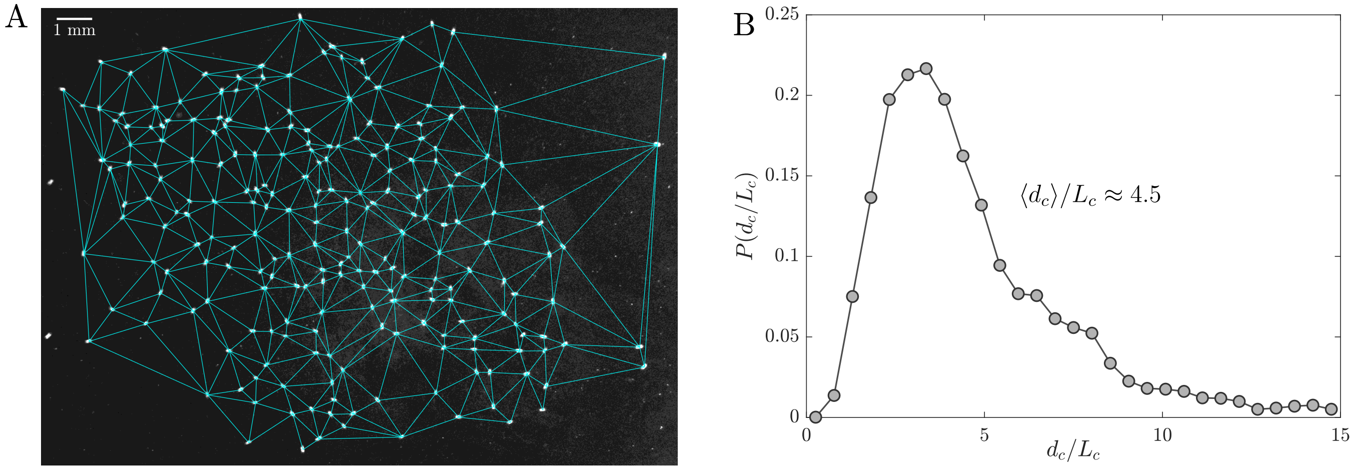


**Fig. 6.** (A) Example of a Delaunay triangulation connecting the feeding cells of a cluster (i.e. the cells for which $v_c < 500\,\mu$m/s). (B) Probability distribution function (PDF) of the length of the straight lines connecting the cells through the Delaunay triangulation $P(d_c/L_c)$, for the feeding cluster presented in Fig. 2A(ii) in the main text.

### 9. Characterization of the collective flow

**A. Experimental method.** The preparation of the observation chamber and cells follows the same procedure as previously described to measure the flow around an isolated cell, except that the cell density inside the chamber is significantly increased. A Hamamatsu ORCA fusion BT camera mounted on an inverted Olympus IX 81 microscope and paired with a 4x objective associated with a magnification changer of 1.6x (i.e., 6.4x magnification in total) provides a fine resolution of $1\,\mu$m/pixel and a field of view of $2.3\,\mathrm{mm} \times 2.3\,\mathrm{mm}$. 8-bit images are captured at 23 fps, with an exposure time of 5 ms. The focal plane is set at $z_{\mathrm{fp}} = 150\,\mu$m. Among cells, a solution consisting of $50\,\mu$l Volvic water seeded with $2\,\mu$m in diameter microbeads is injected, mixed with $50\,\mu$l of a *Chlorogonium sp.* culture. A homogeneous distribution of microbeads is achieved within minutes, thanks to the efficient mixing ability of the feeding cluster. From each image, the Eulerian velocity field is extracted using the PIVlab Matlab program, with a spatial resolution of $40\,\mu$m between adjacent velocity points. To reduce noise and measurement errors, each velocity field is averaged over 5 consecutive frames. Finally, instantaneous streamlines are obtained in each image by solving the advection equation $\dot{\boldsymbol{x}} = \boldsymbol{v}(\boldsymbol{x})$ using the Matlab function ode45. The integration is perfomed from multiple initial positions $\boldsymbol{x}_0$ of material elements arranged on a regular grid, with $\boldsymbol{v}$ the collective velocity field relative to each image. Examples of these streamlines can be seen in Fig. 4C of the main text and in Fig. 7B-D.

**B. Analytical skin friction field on the wall $z = 0$ generated by a single feeding flow, and classification of critical points.** The skin friction field (wall shear) $\boldsymbol{\tau}(x,y)$ imparted by a velocity field $\boldsymbol{v}(x,y)$ on the wall $z = 0$ is defined as

$$\boldsymbol{\tau} = \left.\frac{\partial \boldsymbol{v}}{\partial z}\right|_{z=0}. \tag{13}$$

For $\boldsymbol{v}$ the velocity field generated by a Stokeslet at position $\boldsymbol{x}_f = (x_f, y_f, h)$ and orientation $\boldsymbol{F}_p = (F_{px}, F_{py}, F_{pz})$ over the no-slip boundary $z = 0$ (as described by equation (9)), the $(x,y)$ components of the skin friction field $\boldsymbol{\tau}$ simplify to:

$$\tau_x(x,y) = \frac{3h(-F_z h x_r + F_y x_r y_r + F_x(x_r^2 + \delta^2))}{2\pi(r^2 + h^2 + \delta^2)^{5/2}}, \tag{14}$$

$$\tau_y(x,y) = \frac{3h(-F_z h y_r + F_x x_r y_r + F_y(y_r^2 + \delta^2))}{2\pi(r^2 + h^2 + \delta^2)^{5/2}}, \tag{15}$$

with $(x_r, y_r) = (x - x_f, y - y_f)$, and $r^2 = x_r^2 + y_r^2$.

Critical points $\boldsymbol{x}_p$ of the skin friction field are defined by $\boldsymbol{\tau}(\boldsymbol{x}_p) = 0$, provided that these points are non-degenerate, i.e. $\det(\mathcal{A}) \neq 0$. Critical points are classified either as saddle $\boldsymbol{x}_p^S$, nodes $\boldsymbol{x}_p^N$, or spiral $\boldsymbol{x}_p^V$ according to the sign of the eigenvalues of the linearized skin friction tensor $\mathcal{A}$, which writes:

$$\mathcal{A} = \left.\frac{\partial \boldsymbol{\tau}}{\partial \boldsymbol{x}}\right|_{\boldsymbol{x}=\boldsymbol{x}_p}, \tag{16}$$

with its components

$$\mathcal{A}_{xx} = \frac{\partial \tau_x}{\partial x} = \frac{3h(-F_z h + 2F_x x_r + F_y y_r)}{2\pi(r^2 + h^2 + \delta^2)^{5/2}} - \frac{15 h x_r(-F_z h x_r + F_y x_r y + F_x(x_r^2 + \delta^2))}{2\pi(r^2 + h^2 + \delta^2)^{7/2}},$$

$$\mathcal{A}_{yx} = \frac{\partial \tau_y}{\partial x} = \frac{3F_x h y_r}{2\pi(r^2 + h^2 + \delta^2)^{5/2}} - \frac{15 h x_r(-F_z h y_r + F_x x_r y_r + F_y(y_r^2 + \delta^2))}{2\pi(r^2 + h^2 + \delta^2)^{7/2}},$$

$$\mathcal{A}_{xy} = \frac{\partial \tau_x}{\partial y} = \frac{3F_y h x_r}{2\pi(r^2 + h^2 + \delta^2)^{5/2}} - \frac{15 h y_r(-F_z h x_r + F_y x_r y_r + F_x(x_r^2 + \delta^2))}{2\pi(r^2 + h^2 + \delta^2)^{7/2}},$$

$$\mathcal{A}_{yy} = \frac{\partial \tau_y}{\partial y} = \frac{3h(-F_z h + F_x x_r + 2F_y y_r)}{2\pi(r^2 + h^2 + \delta^2)^{5/2}} - \frac{15 h y_r(-F_z h y_r + F_x x_r y_r + F_y(y_r^2 + \delta^2))}{2\pi(r^2 + h^2 + \delta^2)^{7/2}}.$$

Critical points are nondegenerate points, that is, $\det(\mathcal{A}) \neq 0$.

**C. Strong hyperbolicity of the critical lines.** According to Surana et al. (8), one of the requirements of a 'critical' skin friction line is that it must be strongly hyperbolic (i.e., they exponentially stretch or compress the neighboring skin friction lines), so that a separation or reattachment surface emanate from $\nabla_\perp \cdot \boldsymbol{\tau} < 0$ or $\nabla_\perp \cdot \boldsymbol{\tau} > 0$ all along critical lines, respectively, with $\nabla_\perp \equiv (\partial x, \partial y, 0)$.

In practice, in our system, we observe that most of the 2D manifolds emanating from the saddle points of the skin friction field are not strictly separation/reattachment surfaces, in the sense that they do not consistently exhibit the required saddle-spiral connection and strong hyperbolicity ($|\nabla_\perp \cdot \boldsymbol{\tau}| > 0$) all along the line, as shown in Fig. 7A-B. In fact, some 1D manifolds exhibit

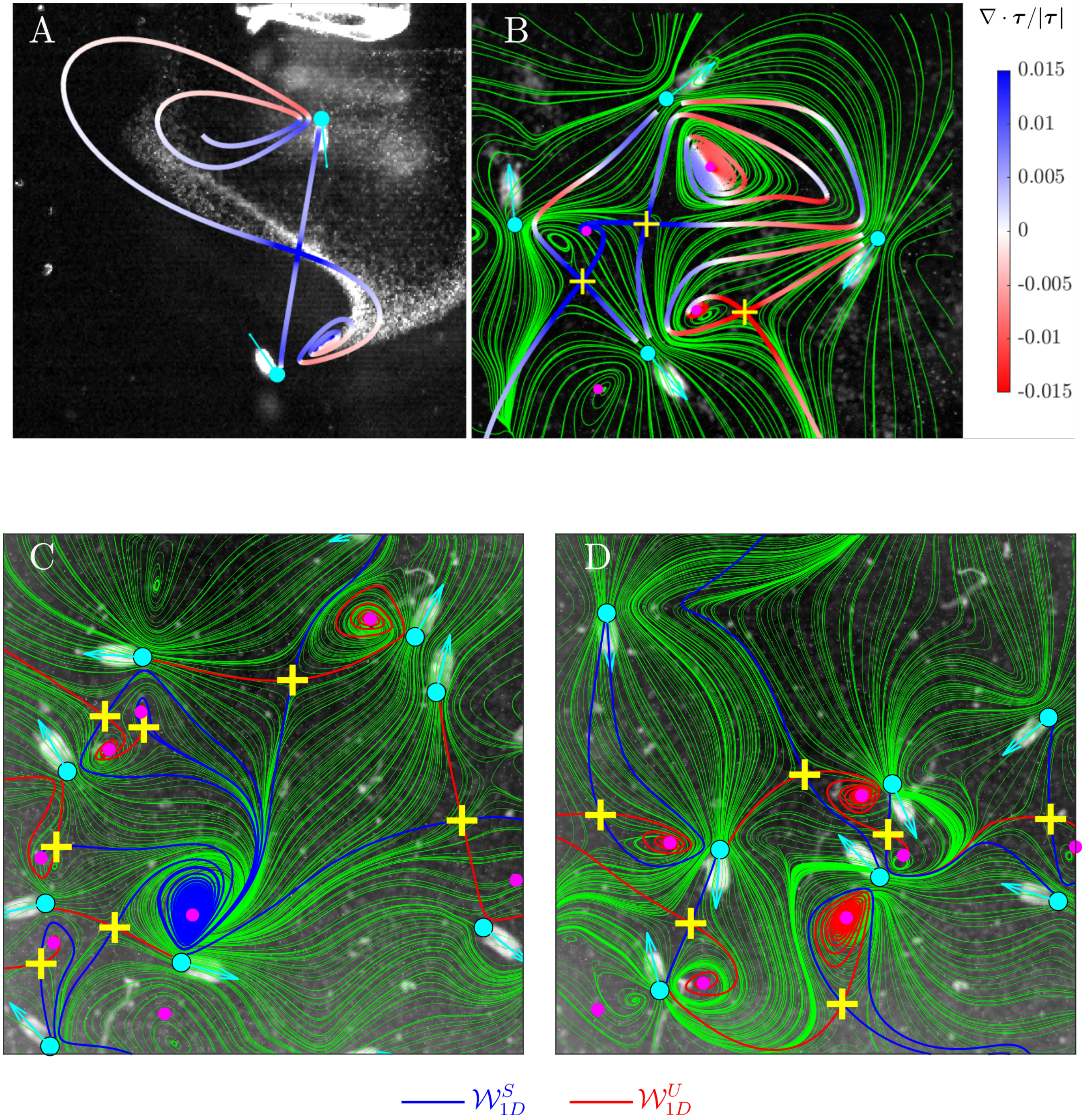


**Fig. 7.** (A) Spatial evolution of $(\nabla_\perp \cdot \boldsymbol{\tau})/|\boldsymbol{\tau}|$ along the 1D manifolds $\mathcal{W}_{1D}^{S}$ and $\mathcal{W}_{1D}^{U}$ of Fig. 4A of the the main text. (B) Spatial evolution of $(\nabla_\perp \cdot \boldsymbol{\tau})/|\boldsymbol{\tau}|$ along the 1D manifolds $\mathcal{W}_{1D}^{S}$ and $\mathcal{W}_{1D}^{U}$ for another configuration of feeding cells, superimposed to the experimental flow (in green). The feeding forces locations and orientations are represented by cyan disk and arrow respectively. The colorbar refers only to the two panels (A) and (B). (C-D) Two other examples of feeding cells configurations, with the analytically computed $\mathcal{W}_{1D}^{S}$ (blue lines) and $\mathcal{W}_{1D}^{U}$ (red lines), superimposed on the experimentally measured flow in green streamlines. In all cases, the experimental velocity field was measure in the focal plane $z_{\mathrm{fp}} = 150\,\mu$m above the no-slip wall.

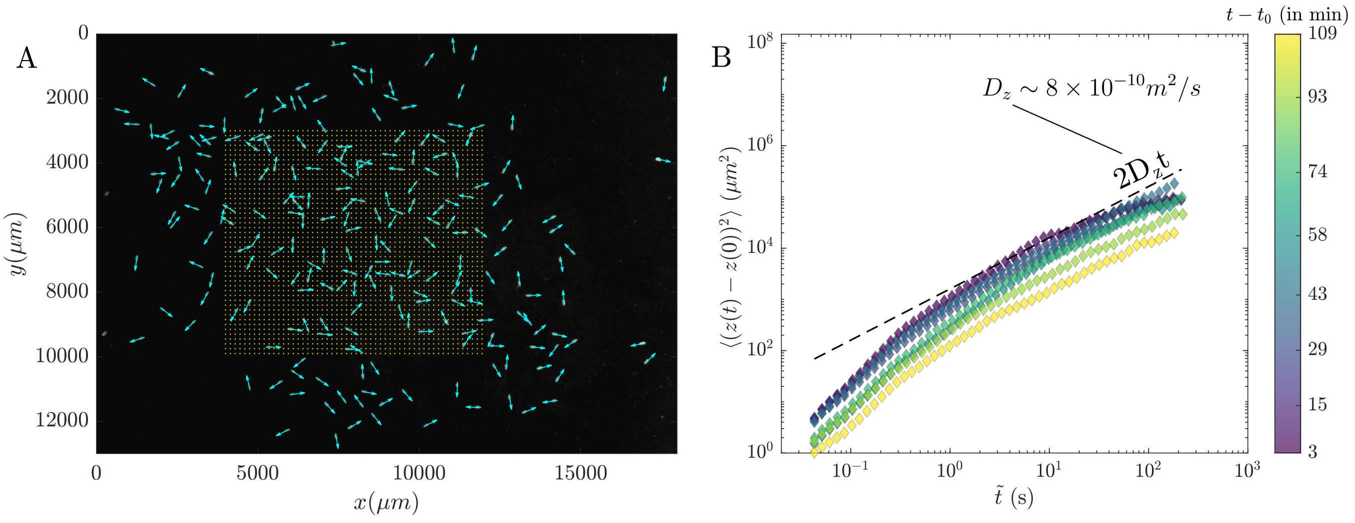


**Fig. 8.** (A) Picture of the initial positions $\boldsymbol{x}_0$ (seen as yellow points) of 2538 fluid elements arranged on a regular square grid, and set at a constant altitude $z = 150\,\mu$m. The grid of tracers covers most of the feeding cluster (here for the first image of movie $t - t_0 = 15$ min). Volume force positions and orientations are depicted as cyan dots and arrows on the image, and are set by the cells positions and orientations, respectively. (B) Temporal evolution of the variance of displacement (MSD) of the fluid particles in the $z$-direction $\sigma_z^2(t)$.

a sign reversal of $\nabla_\perp \cdot \boldsymbol{\tau}$ along their length. This behavior is expected given that the flow in our system is 'open': the flow intensity vanishes with distance $z$ from the wall, implying that the influence of the 2D manifolds diminishes as we move farther from the boundary. However, as long as we remain sufficiently close to the solid boundary $z = 0$, the manifolds retain their role in shaping and governing the flow. In fact, as demonstrated in Fig. 7C-D, regardless of their hyperbolicity, the alignment between the theoretical 1D manifolds and the experimental flow is nearly perfect, even as the number of cells increases. Thus, these manifolds, whether or not they possess strong hyperbolicity, remain relevant in defining the flow structure near the wall and contributing to the flow skeleton and Lagrangian Coherent Structures (LCSs).

### 10. Fluid dispersion

**A. Numerical quantification of the fluid dispersion coefficient.** We quantify the temporal evolution of the mean squared displacements (MSDs) in both the $xy$-plane and the $z$-direction of 2538 fluid particles numerically advected by the collective velocity field $\dot{\boldsymbol{x}} = \boldsymbol{v}(\boldsymbol{x}(t), t_j)$, with $\boldsymbol{v}$ the analytical velocity field updated in each image $j$. The numerical integration of the advection equation is carried out using the ode45 Matlab solver to precision $10^{-7}$. The tracer trajectories are integrated over the 3.5 min duration of each movie, for each movie presented in Fig. 5C of the main text, i.e. from $t - t_0 = 3$ min to $t - t_0 = 109$ min. The initial tracer positions $\boldsymbol{x}_0$ are arranged on a regular square grid that covers most of the feeding cluster in the $xy$-plane, with an initial altitude of $z_0 = 150\,\mu$m (see Fig. 8A). This grid is consistent across all movies from $t - t_0 = 15$ min to $t - t_0 = 109$ min, but for the first movie $t - t_0 = 3$ min, the grid $\boldsymbol{x}_0$ is much smaller to accommodate the still-emerging feeding cluster.
As shown in Fig. 8B, the MSDs in the $z$-direction $\sigma_z^2(t)$ for the latest movies reach a Fickian regime, while the two earliest movies ($t - t_0 = 3$ min and $t - t_0 = 15$ min) are subdiffusive. This is expected because, at these early times, the cells are more stable and tightly packed, which causes the fluid particles to travel farther upward in the $z$ direction (possibly along the 2D unstable manifolds $\mathcal{W}_{2D}^U$), where the magnitude of average velocity fluctuations decays strongly. The maximum transverse fluid dispersion coefficient observed, although briefly, is $D_z^{\text{fl}} = 8 \times 10^{-10}\,\text{m}^2$/s.

**B. Estimation of the fluid dispersion coefficient by a suspension of swimmers of equivalent population density moving in an unbounded fluid.** We estimate the effective fluid diffusivity induced by a group of *S. lemnae* cells crawling, using the model of Lin et al. (9), which predicts the effective diffusivity $\kappa$ from the fluid displacement due to swimming bodies (Darwin drift) in an unbounded fluid under low Reynolds number conditions. In this model, the suspension of swimmers is assumed to be dilute, which means that there are no hydrodynamical interactions between individual fluid disturbances. The swimmers are modeled as squirmers, which are spheres in Stokes flow with a velocity prescribed on their surface representing the effect of the continuous beating of cilia. In their model, the major contribution to fluid particle displacement is due to the far-field stresslet term of the velocity field induced by the squirmer, leading to the following equation for the effective diffusivity $\kappa$:

$$\kappa \approx 2.1\beta^2 U n R^4, \tag{17}$$

with $R$ the radius of the axially symmetric swimmer, $U$ the constant speed over the full path length $\lambda$, $n$ the number density of swimmers, and $\beta$ is the stresslet coefficient (or squirmer parameter), which is a free parameter in their model. To calculate

$\beta$ for our case, we express it as a function of the stresslet strength $p$. Indeed, the far-field flow around a crawling *S. lemnae* at $U = v_c^{\text{free}} = 770\,\mu$m/s can be approximated as a stresslet, defined as:

$$\boldsymbol{v}(\boldsymbol{r}) = \frac{p}{|\boldsymbol{r}|^2}[3(\hat{\boldsymbol{r}}\cdot\hat{\boldsymbol{e}})^2 - 1]\hat{\boldsymbol{r}}, \quad p = -\frac{lF}{8\pi\mu}, \tag{18}$$

with $\hat{\boldsymbol{r}} = \boldsymbol{r}/|\boldsymbol{r}|$, $\boldsymbol{r}$ the distance vector relative to the stresslet, $\hat{\boldsymbol{e}}$ the swimming direction, $l$ the distance between the two opposite forces composing the stresslet, $F$ the force magnitude and $\mu$ the fluid viscosity. We use the values $l \approx \delta = 43\,\mu$m, $F = |\boldsymbol{F}_p| = 1200$ pN, $\mu = 10^{-3}$ N s/m$^2$, to obtain the stresslet strength $p = 2.05\times10^6\,\mu$m$^3$/s.
Next, we seek the relationship between $p$ and $\beta$ by starting with the flow field generated by an axisymmetric squirmer of radius $R$, expressed in the comoving cell frame as (10, 11):

$$v_r(r,\theta) = \frac{2}{3}\left(\frac{R^3}{r^3} - 1\right)B_1P_1(\cos\theta) + \sum_{n=2}^{\infty}\left(\frac{R^{n+2}}{r^{n+2}} - \frac{R^n}{r^n}\right)B_nP_n(\cos\theta), \tag{19}$$

$$v_\theta(r,\theta) = \frac{2}{3}\left(\frac{R^3}{2r^3} + 1\right)B_1V_1(\cos\theta) + \sum_{n=2}^{\infty}\frac{1}{2}\left(n\frac{R^{n+2}}{r^{n+2}} + (2-n)\frac{R^n}{r^n}\right)B_nV_n(\cos\theta), \tag{20}$$

with $\theta$ the angle with the swimming axis $\hat{\boldsymbol{e}}$. The first mode $n = 1$ corresponds to the source term $\boldsymbol{v}^{\text{s}}$, and the second mode $n = 2$ to the stresslet term $\boldsymbol{v}^{\text{str}}$. The squirmer parameter is $\beta = B_2/|B_1|$, and the swimming speed $U = 2B_1/3$. On the swimming axis $\hat{\boldsymbol{e}}$, $\theta = 0$, $v_\theta(r,\theta) = 0$, and the stresslet term in equation (19) is:

$$v_r^{\text{str}}(r,\theta) = \left(\frac{R^4}{r^4} - \frac{R^2}{r^2}\right)B_2P_2(\cos\theta). \tag{21}$$

Since we have $B_2 = B_1\beta = \frac{3}{2}U\beta$ and $P_2(\cos\theta) = \frac{1}{2}(3\cos^2\theta - 1) = 1$ for $\theta = 0$, the previous equation becomes:

$$v_r^{\text{str}}(r,0) = \frac{3}{4}U\beta\left(\frac{R^4}{r^4} - \frac{R^2}{r^2}\right). \tag{22}$$

In the far-field flow $r \gg R$, we approximate $(R^4/r^4 - R^2/r^2) \sim -R^2r^2$. We have finally:

$$v_r^{\text{str}}(r,0) = -\frac{3\beta U}{4}\frac{R^2}{r^2}. \tag{23}$$

By comparing this with equation (18), we deduce the following relation between stresslet strength $p$ and squirmer parameter $\beta$:

$$\beta = -\frac{4p}{3UR^2}. \tag{24}$$

Taking the values $U = v_c^{\text{free}} = 770\,\mu$m/s and $R = \delta = 43\,\mu$m, we get $\beta = 1.92$. Substituting this value of $\beta$ (from equation (24)) into equation (17), we obtain the final expression for the effective diffusivity $\kappa$:

$$\kappa \approx \frac{3.7p^2n}{U}. \tag{25}$$

We estimate the number density of swimmers in the feeding cluster of movie $t - t_0 = 15$ min to be $n \approx 220\,\text{cells}/22\,\text{mm}^3 = 10^{-8}\,\text{cells}/\mu\text{m}^3$. Substituting these values into equation (25), we predict the effective fluid diffusivity due to crawling *S. lemnae* cells to be approximately $\kappa \approx 2.10^{-10}\,\text{m}^2/\text{s}$.

### 11. Definition of the mixing region

We define the mixing region as the volume that encompasses most of the collective flow generated by the feeding cluster. Fluid particles crossing the boundaries of this region are no longer considered in the calculation of the Finite-Time Lyapunov Exponents (FTLEs). The mixing region is bounded in the $xy$-plane by the outer edge of the feeding cells' cluster and by a maximum altitude $z_{\text{max}}$. The boundary of the mixing region in the $xy$-plane is computed using the Matlab function 'boundary' applied to the set of feeding cells (cells for which $v_c < v_c^{\text{max}}$). This boundary is updated at each image. An example of such a boundary is shown in Fig. 9A.
The maximum altitude that limits the mixing region in the $z$-direction is set to $z_{\text{max}} = 20h$. Given the fluid diffusivity in the $z$-direction $D_z^{\text{fl}} = 8\times10^{-10}\,\text{m}^2/\text{s}$, the average time for fluid particles to reach the altitude $z = z_{\text{max}}$ is $t \sim (z_{\text{max}} - z_0)^2/D_z^{\text{fl}} \approx 900$ s, which is much longer than the 3.5-minute duration of a typical movie. As shown in Fig. 9B, only 8% of the fluid particles cross the $z_{max}$ threshold after $t = 3.5$ min.

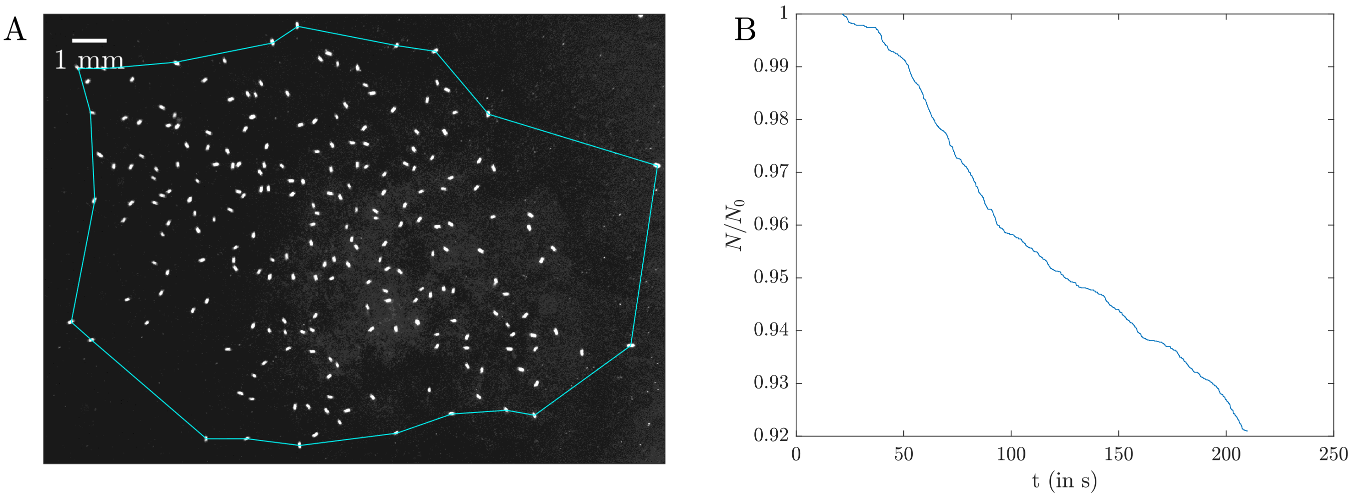


**Fig. 9.** (A) Extent of the mixing region in the $xy$-plane delimiting the cluster of feeding cells, obtained with the Matlab function 'boundary'. (B) Loss of fluid particles in time by crossing the fixed altitude limit $z_{\text{max}} = 20h$, over the duration of the movie $t - t_0 = 15$ min.

### 12. Computation of the finite-time Lyapunov exponents (FTLEs), and their average $\lambda$

**A. Method.** We compute the finite-time Lyapunov exponents (FTLEs) $\tilde{\mu}(t)$, which measure the exponential rate of fluid stretching, as follows: in a first step, we compute the Lagrangian trajectories $\mathbf{x}(t)$ by solving the advection equation

$$\frac{d\mathbf{x}(\mathbf{X},t)}{dt} = \boldsymbol{v}(\mathbf{x}(t),t), \tag{26}$$

starting at $t = 0$ from $24 \times 27 = 648$ different initial positions $\mathbf{X}$ arranged on a regular square grid fitting most of the feeding cluster and set at $z = 150\,\mu$m distance from the boundary. Here, $\boldsymbol{v}$ is the collective fluid velocity field made of the sum of the individual pumping flows $\boldsymbol{v}_i$ described by equation (9):

$$\boldsymbol{v}(\mathbf{x},t_j) = \sum_{i=1}^{N} \boldsymbol{v}_i(\mathbf{x},t_j), \tag{27}$$

where $\boldsymbol{v}(t_j)$ is updated at each image $j$ based on the evolving cell positions and orientations, i.e. at regular times $t_j = j\Delta t$ with $\Delta t = 0.043$ s the period between two consecutive images. An important point here is that any noise artifacts inevitably arising from small errors in image cell detection will impart large errors in the calculation of the tensor of the deformation gradient $\mathbf{F}$ close to the volume forces. Therefore, we enforce the immobility of the cells as long as $v_c < v_c^{\text{max}} = 500\,\mu$m/s, keeping their positions and orientations constant during that time. This might introduce a slight discrepancy between the true positions of cells and their modeled force locations, but should have a minimal impact on the overall fluid deformation measurements, as cells generally crawl faster than $v_c^{\text{max}}$. The advection equation (26) is solved using the ode45 Matlab solver (Runge-Kutta method) to precision $10^{-8}$. For the average of the FTLEs to converge towards the constant value $\lambda$, the trajectories are integrated over two full movie durations in a loop: when we reach the end of a movie, the trajectory is continued from its last position. As we assume that *S. lemnae* average behavior is stationary, this does not induce errors on the average fluid stretching measurements. The few trajectories crossing out the limits of the mixing region are stopped.

In a second step, we compute the deformation gradient tensor $\mathbf{F}(t)$ along the Lagrangian trajectories, defined as $\mathbf{F}(t) = \text{d}\mathbf{x}(t)/\text{d}\mathbf{X}$ where $\text{d}\mathbf{x}(t)$ and $\text{d}\mathbf{X}$ denote, respectively, material vectors in the Eulerian and Lagrangian frames. Thus $\mathbf{F}(t)$ characterizes the deformation from the infinitesimal reference vector $d\mathbf{X} = d\mathbf{x}(t = 0, \mathbf{X})$ to $d\mathbf{x}$(t), as $d\mathbf{x} = \mathbf{F} \cdot d\mathbf{X}$. From this definition, $\mathbf{F}(t)$ evolves along a Lagrangian trajectory according to the differential equation:

$$\frac{d\mathbf{F}(t)}{dt} = \boldsymbol{\epsilon}(t)\mathbf{F}(t), \quad \mathbf{F}(0) = \mathbf{I}, \tag{28}$$

with $\boldsymbol{\epsilon}(t) = \nabla\boldsymbol{v}(\mathbf{x}(t),t)^\top$ the velocity gradient tensor and $\mathbf{I}$ the identity tensor taken as initial conditions. The components of $\boldsymbol{\epsilon}(t)$ are calculated analytically from the individual pumping flow of equation (9). Finally, the FTLE along each trajectory $\tilde{\mu}(t)$ is computed from the largest eigenvalue $\nu(t)$ of the Cauchy-Green tensor $\mathbf{B}(t) = \mathbf{F}(t)\mathbf{F}^\top(t)$ as:

$$\tilde{\mu}(t) = \frac{1}{2t}\ln\nu(t). \tag{29}$$

However, for large deformation (especially near volume forces), numerical errors can blow up exponentially. To mitigate this, we employ the discrete QR factorization method, which decomposes at regular time steps the fundamental solution matrix $\mathbf{F}$

into two matrices $\mathbf{F} = \mathbf{QR}$, where $\mathbf{Q}$ is an orthogonal matrix and $\mathbf{R}$ an upper triangular matrix. We follow the procedure described by (12): starting from the initial conditions $\mathbf{Y}(0) = \mathbf{Q}_0 = \mathbf{I}$, we solve at regular time steps $t_j = j\Delta t$ (with $\Delta t$ the period between successive images $j$) the equation for $\mathbf{Y}(t)$:

$$\frac{d\mathbf{Y}(t)}{dt} = \boldsymbol{\epsilon}(t)\mathbf{Y}(t),\ \mathbf{Y}(t_j) = \mathbf{Q}_j \quad t_j \le t \le t_{j+1}, \tag{30}$$

then decompose the solution at $t_{j+1}$ into $\mathbf{Y}(t_{j+1}) = \mathbf{Q}_{j+1}\mathbf{R}_{j+1}$. The differential equation (30) is solved in a loop for $j$ using an explicit Runge-Kutta ode45 Matlab solver with precision $10^{-8}$. The deformation gradient tensor at time $t_{j+1}$ is finally:

$$\mathbf{F}_{j+1} = \mathbf{Q}_{j+1}\mathbf{R}_{j+1}\mathbf{R}_j...\mathbf{R}_1 = \mathbf{Q}_{j+1}\prod_{k=j+1}^{1}\mathbf{R}_k. \tag{31}$$

We keep track of the numerical errors by checking that $\det(\mathbf{F}) - 1 = 0$ (which for incompressible flow should remain valid). The three FTLEs along each trajectory $i$ are defined as:

$$\mu_{i,\alpha}(t_j) = \frac{1}{t_j}\ln||\prod_{k=j}^{1}(\mathbf{R}_{k,i})_{\alpha\alpha}||,\ \alpha = 1,\ 2,\ 3, \tag{32}$$

from which we extract the largest FTLE along each trajectory as $\tilde{\mu}_i = \max_{\alpha}\left(\mu_{i,\alpha}\right)$. Finally, since the video duration is finite, we use the ergodic property of random chaotic flows to calculate the Lyapunov exponent $\lambda$ as the ensemble average of $\tilde{\mu}_i$ over all trajectories $i$:

$$\lambda = \left\langle \tilde{\mu}_i(T) \right\rangle, \tag{33}$$

with $T$ the total integration time equal to twice the duration of a movie (approximately about 420 s).

**B. Validation of the method for measuring fluid deformations for the steady pumping flow case.** Our method for measuring infinitesimal deformation along trajectories, combining the analytical development of $\boldsymbol{\epsilon}(t) = \nabla \boldsymbol{v}(t)^\top$ and the numerical estimation of the fluid trajectory $\mathbf{x}(t)$ and $\mathbf{F}(t)$, is validated for the simple steady pumping flow described by equation (9), where the pumping force $\boldsymbol{F}_p$ is parallel to the no-slip boundary, i.e. $\phi = 0$. In this case, we demonstrate that the method can recover the time-reversal symmetry of the transverse deformation kinematics relative to the position of the volume force. Fig. 10A shows a particle fluid trajectory $\mathbf{x}(t)$ advected by the flow from the point $(2.5, 0, 1)L_c$ over a complete cycle $t_{\text{cycle}}$. The trajectory exhibits perfect symmetry with respect to $x = 0$, with the particle traveling an equal distance on both sides of the force. The components of the velocity gradient tensor $\nabla \boldsymbol{v}(\mathbf{x}(t))$ calculated along the trajectory are displayed on Fig. 10B, with peaks corresponding to the moment when the particle is closest to the volume force. For that specific case, we rewrite the velocity gradient tensor $\boldsymbol{\epsilon}(t) = \nabla \boldsymbol{v}(t)^\top$ from the Cartesian coordinate system$\{x, y, z\}$ to the streamline coordinate system $\{x', y', z'\}$, see (13), where the new frame axis $x'$ is continuously reoriented along the streamline, while the other two axes $y'$ and $z'$ are continuously orthogonal to the streamline. In this way $\boldsymbol{\epsilon}'(t)$ - and so $\mathbf{F}'(t)$ through equation (28) - are made upper-triangular, and the diagonal components of $\mathbf{F}'(t)$ are then directly representative of the stretching experienced by the particle along the axis $\{x', y', z'\}$. These components $F'_{ii}$ are shown on Fig. 10C, and demonstrate that there is no net deformation after one complete cycle along the streamline direction $x'$ or orthogonal to the streamline. This behavior is consistent with the expected time-reversal symmetry of the flow transverse to the streamline. However, note that the total net deformation after one cycle does not cancel strictly, as shown by the eigenvalues of the Cauchy-Green tensor $\mathbf{B}$ in Fig. 10D, which reveal a net deformation $\approx 2$, which originates entirely from shear deformation parallel to the streamline.

### 13. Second yeast injection experiment and anomalous fluid transport

**A. Experimental method.** The second localized yeast injection is performed approximately 2 h after a first injection which triggered the development of a feeding cell cluster. The yeast is introduced near the previously established cluster. High-speed imaging is then performed over 977 s at 23 fps with a Hamamatsu ORCA fusion BT camera, mounted on a stereo Olympus SZX16 microscope with a 0.7x objective. This setup provides a resolution of $9.6\,\mu$m/pixel and a field of view of 22.1 mm × 22.1 mm. The exposure time is set to 10 ms.

**B. Method for the numerical computation of the tracer MSD.** Initially, 1000 material elements are randomly positioned within the yeast blob detected in the first image, at a common height of $z = 150\mu$m, to represent the transport of yeast particles. They are numerically advected by solving the advection equation over the collective flow generated by the established feeding cluster, up to time $t = 976.8$s (see Fig. 11B). The mean squared displacement (MSD) is computed from the tracer trajectories. Unlike the previous tracer MSDs calculated in section 3A for the first yeast injection, as the yeast in this case is pumped towards

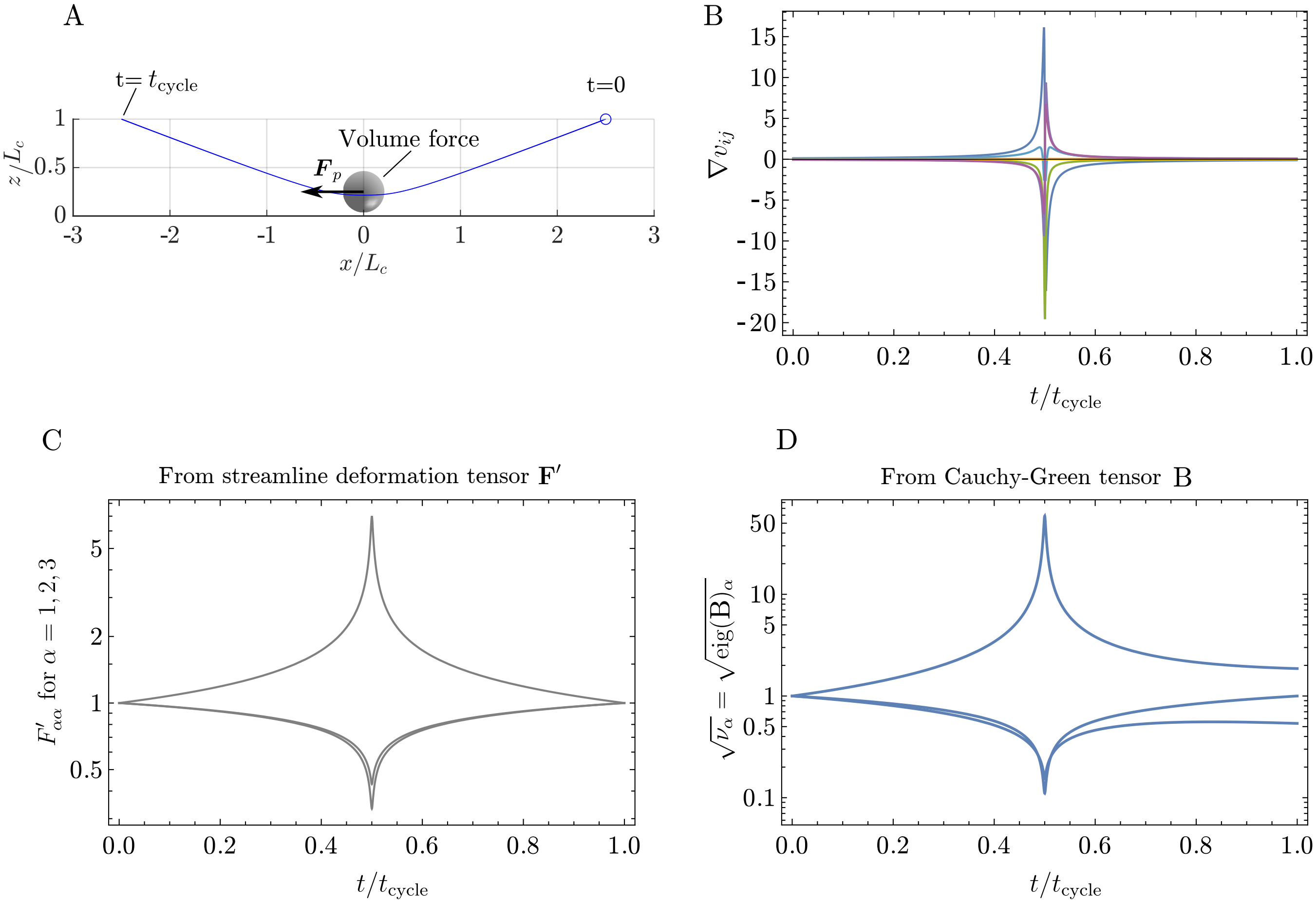


**Fig. 10.** (A) Streamline $\mathbf{x}$ of a particle fluid advected by a steady volume force over the no-slip boundary $z = 0$, with the force direction $\boldsymbol{F}_p$ parallel to the boundary. (B) Temporal evolution of the nine analytical components of $\nabla \boldsymbol{v}(\mathbf{x}(t))$ for the streamline in (A). (C) The three diagonal components of $\mathbf{F}'$: one indicates the deformation along the streamline, the two others the deformation in the plane perpendicular to the streamline. (D) Eigenvalues of the Cauchy-Green tensor $\mathbf{B} = \mathbf{F}'\mathbf{F}'^{\top} (= \mathbf{F}\mathbf{F}^{\top})$.

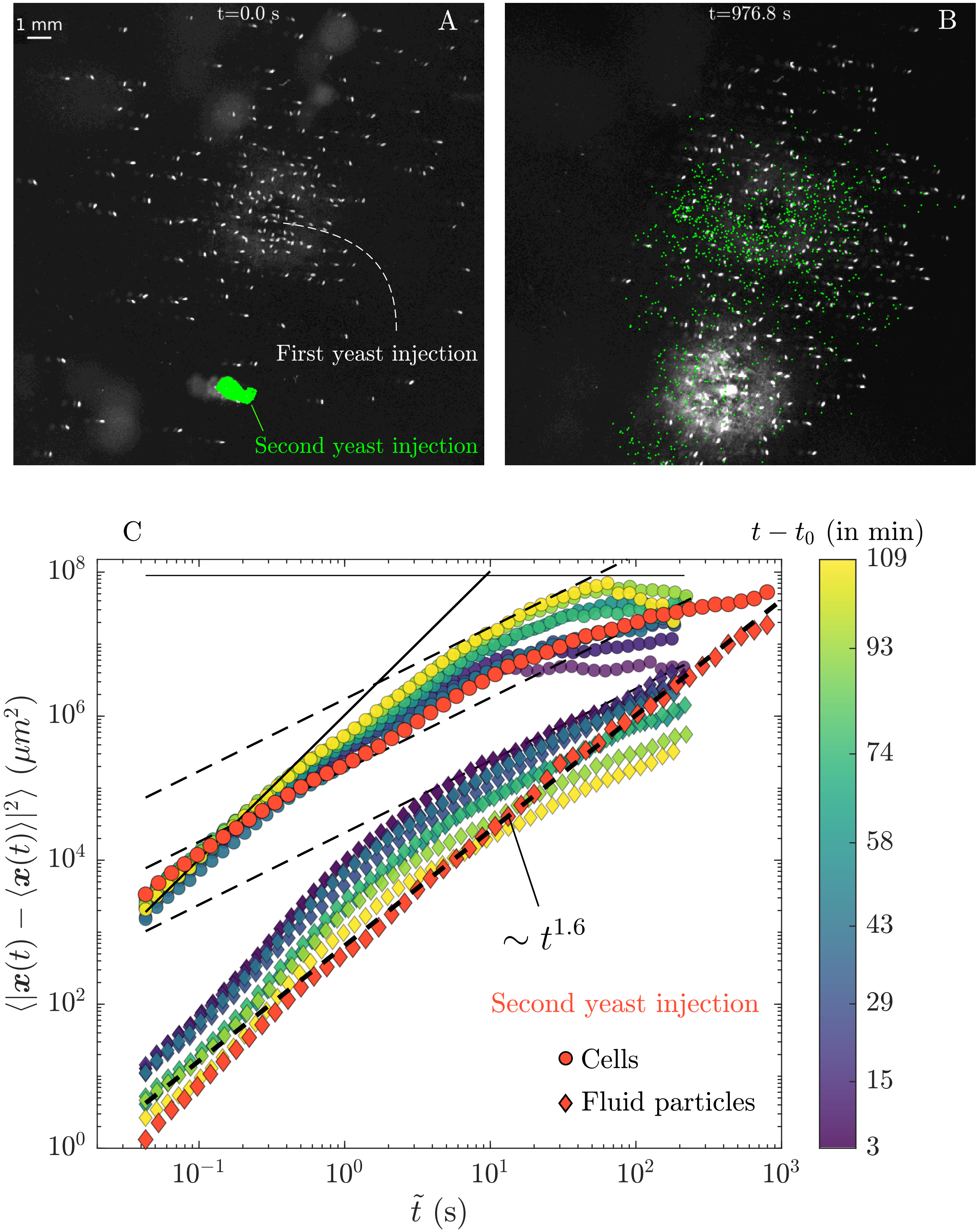


**Fig. 11.** (A) Second localized yeast injection, near an already established feeding cell cluster, triggered by a first injection. 1000 particle tracers (green dots) are initially positioned randomly within the yeast blob, at a constant height $z = 150\mu$m (B) Evolution of the feeding cluster after $t = 977$s. The tracers particles have been advected by the associated collective flow towards the cluster. At this time, the spatial distribution of cells and tracers closely match (C) The MSDs for both the cells and simulated fluid particles are overlaid on top of Fig. 5C from the main text, showing the results for the second yeast injection near the established feeding cluster.

the cluster, the fluid spreading induced by the feeding cluster is anisotropic, and consequently $\langle \boldsymbol{x}(t) \rangle \neq \boldsymbol{x}(0)$. Thus, the MSD is computed based on the displacement variance relative to the average tracer position: $\sigma_x^2(t) = \langle |\boldsymbol{x}(t) - \langle \boldsymbol{x}(t) \rangle|^2 \rangle$. We observe that MSD exhibits an anomalous scaling $\sigma_x^2(t) \sim t^{\alpha}$, with $\alpha = 1.6$. A key point here is the choice of both the initial tracer positions and the starting advection time. Since the tracers are positioned just outside the feeding cluster rather than within it, there is a delay before the cells begin to detect and pump the newly injected yeast blob. This occurs approximately 17 s after the second injection, which is set as the starting time for the tracer advection. Furthermore, we found that altering the initial positions of the tracers slightly affects the anomalous exponent $\alpha$, which fluctuates between $\alpha = 1.4$ and $\alpha = 1.6$ (the latter corresponding to tracers initially placed within the yeast blob). In all cases $\alpha > 1$, provided that the initial positions of the particles are not set too far from the cluster.
